# Restoration of excitation/inhibition balance enhances neuronal signal-to-noise ratio and rescues social deficits in autism-associated *Scn2a*-deficiency

**DOI:** 10.1101/2025.03.04.641498

**Authors:** Jingliang Zhang, Muriel Eaton, Xiaoling Chen, Yuanrui Zhao, Shivam Kant, Brody A. Deming, Kothandaraman Harish, Huynhvi P. Nguyen, Yue Shu, Shirong Lai, Jiaxiang Wu, Zhefu Que, Kyle W. Wettschurack, Zaiyang Zhang, Tiange Xiao, Manasi S. Halurkar, Maria I. Olivero-Acosta, Ye-Eun Yoo, Nadia A. Lanman, Wendy A. Koss, William C. Skarnes, Yang Yang

**Affiliations:** Borch Department of Medicinal Chemistry and Molecular Pharmacology, College of Pharmacy, Purdue University; Purdue Institute for Integrative Neuroscience, Purdue University; Department of Comparative Pathobiology, Purdue University; Purdue University Center for Cancer Research, Purdue University; Department of Basic Medical Sciences, College of Veterinary Medicine, Purdue University; Office of the Executive Vice President for Research and Partnerships, Purdue University; The Jackson Laboratory for Genomic Medicine

**Keywords:** *SCN2A/Scn2a*, voltage-gated sodium channel, autism, sociability, E/I imbalance, signal-to-noise ratio, brain organoid

## Abstract

Social behavior is critical for survival and adaptation, which is profoundly disrupted in autism spectrum disorders (ASD). Social withdrawal due to information overload was often described in ASD, and it was suspected that increased basal noise, i.e., excessive background neuronal activities in the brain could be a disease mechanism. However, experimental test of this hypothesis is limited. Loss-of-function mutations (deficiency) in *SCN2A*, which encodes the voltage-gated sodium channel Na_V_1.2, have been revealed as a leading monogenic cause of profound ASD. Here, we revealed that *Scn2a* deficiency results in robust and multifaceted social impairments in mice. *Scn2a*-deficient neurons displayed an increased excitation-inhibition (E/I) ratio, contributing to elevated basal neuronal noise and diminished signal-to-noise ratio (SNR) during social interactions. Notably, the restoration of *Scn2a* expression in adulthood is able to rescue both SNR and social deficits. By balancing the E/I ratio and reducing basal neuronal firing, an FDA-approved GABA_A_ receptor-positive allosteric modulator improves sociability in *Scn2a*-deficient mice and normalizes neuronal activities in translationally relevant human brain organoids carrying autism-associated *SCN2A* nonsense mutation. Collectively, our findings revealed a critical role of the Na_V_1.2 channel in the regulation of social behaviors, and identified molecular, cellular, and circuitry mechanisms underlying *SCN2A*-associated disorders.

**HIGHLIGHTS:**

1. Na_V_1.2 deficiency leads to pronounced social deficits in mice.
2. Na_V_1.2 deficiency results in an overall enhanced E/I ratio, elevated basal neuronal activity, and impaired signal-to-noise ratio.
3. Both the enhanced E/I ratio and impaired sociability are reversible through the restoration of Na_V_1.2 expression in adulthood.
4. Targeted restoration of Na_V_1.2 in striatum-projecting neurons rescues social impairments.
5. GABA transmission is reduced in both mouse and human organoid models of *SCN2A* deficiency, and acute systemic administration of GABA_A_ receptor-positive allosteric modulators restores sociability.

**Graphical abstract:** 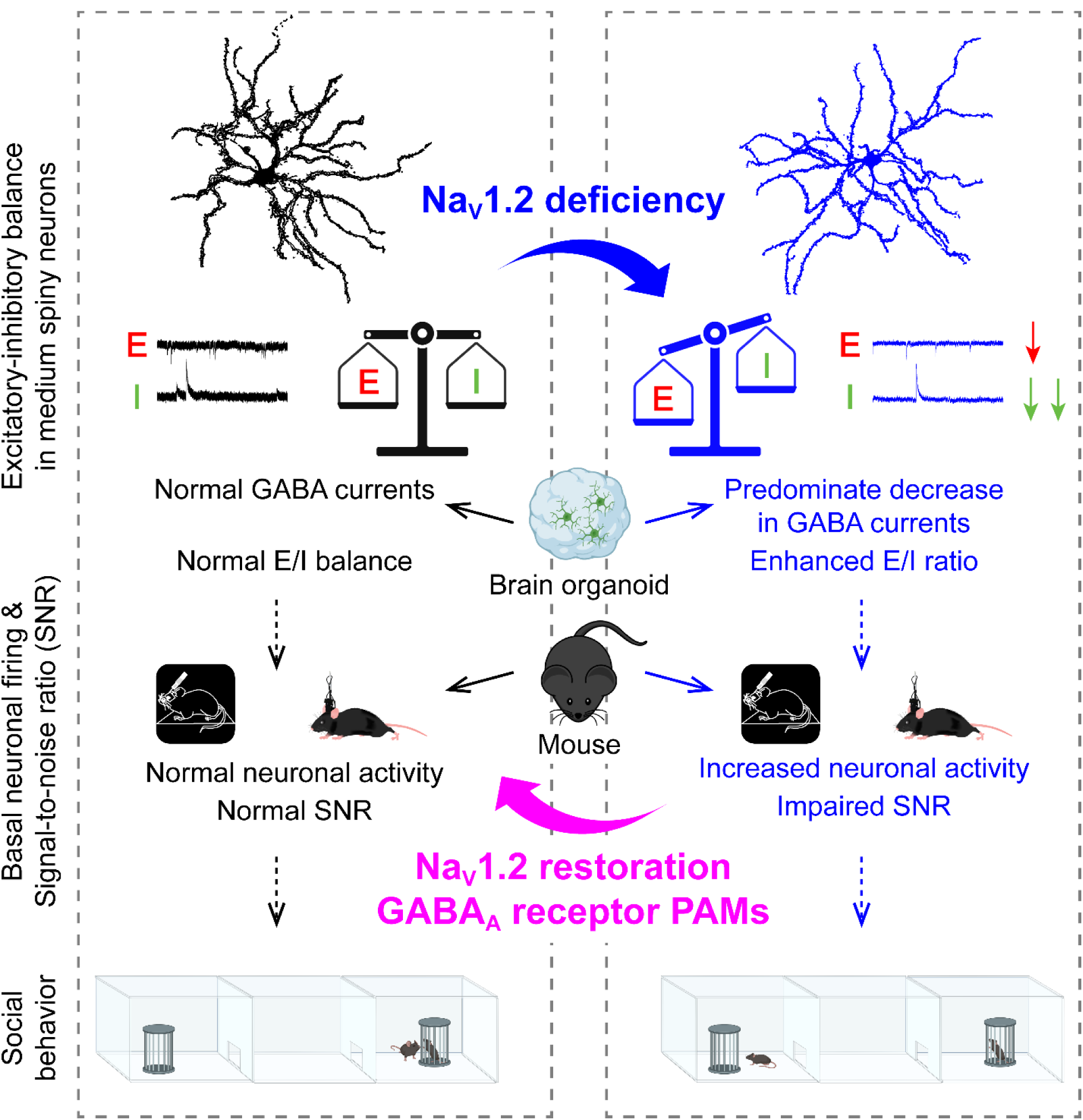

Graphical abstract: Severe *Scn2a* deficiency leads to a predominate decrease in GABA transmission with an overall enhanced E/I ratio, elevated basal neuronal activity, impaired SNR, and social deficits in adult Na_V_1.2-deficient mice.

## INTRODUCTION

The incidence of autism spectrum disorder (ASD) has risen dramatically from 1 in 150 children in 2000^1^ to 1 in 36 in 2020^2^. Although ASD is largely polygenic^3^, mutations in a single gene can also produce severe forms of the disorder. Regardless of its heterogeneous manifestations, impaired social interaction remains a core diagnostic feature of ASD. Recent large-scale genetic studies in humans have identified *SCN2A* as one of the most prominent ASD causal genes^4,5^. *SCN2A* encodes Na_V_1.2, a voltage-gated sodium channel that is critical for action potential generation and synaptic function in the central nervous system^6^. Protein-truncating and loss-of-function variants in *SCN2A*, collectively termed *SCN2A* deficiency, are among the most common genetic alterations associated with ASD. Although *SCN2A* mutations in patients are heterozygous, studies in *Scn2a^+/–^* mice have yielded inconsistent results regarding social behaviors^7–9^. This suggests that a 50% reduction in Na_V_1.2 expression in mice may be insufficient to elicit overt phenotypes, potentially due to compensatory mechanisms^10^. Given the essential role of *SCN2A* in neuronal excitability, complete knockout results in perinatal lethality^11^, suggesting a more severe, yet incomplete, reduction in Na_V_1.2 expression may be necessary to unmask pronounced behavioral deficits relevant to ASD.

We employed a gene-trap (gt) mouse model (*Scn2a^gt/gt^*) that exhibits Na_V_1.2 deficiency yet survives to adulthood^12^. Na_V_1.2 is widely expressed in brain regions critical for social behavior, including the striatum^13,14^, making this model well-suited for investigating the neurobiological basis of social impairments associated with *Scn2a* deficiency. Recent theories propose that an imbalance in excitatory and inhibitory (E/I) synaptic transmission is central to ASD^15–17^, although it remains unclear whether such imbalances directly cause impaired behavioral function. Altered neuronal signal-to-noise ratio (SNR) during social interactions has also been highlighted^18^. However, the mechanisms underlying SNR determination remain ambiguous. Investigating whether E/I imbalances critically affect the processing of social signals may therefore provide valuable insights into potential interventions for social impairments. Importantly, the gene-trap strategy not only results in a marked reduction in *Scn2a* expression but also enables flexible genetic manipulations to evaluate the bidirectional role of *Scn2a* in determining SNR and regulating social behaviors.

In the present study, we systematically examined the behavioral, electrophysiological, and neurophysiological consequences of Na_V_1.2 deficiency. We assessed social behavior in *Scn2a^gt/gt^* mice using multiple assays, including the three-chamber sociability, reciprocal interaction, and resident-intruder tests. To evaluate synaptic transmission, we conducted patch-clamp recordings from medium spiny neurons in the nucleus accumbens, while high-density Neuropixels recordings and *in vivo* calcium imaging provided insights into the basal and behaviorally evoked neuronal activity in the striatum. Additionally, we investigated whether global or regionally selective restoration of *Scn2a* expression in adulthood can rescue the observed deficits. Guided by electrophysiological assessments of inhibitory transmission and RNA sequencing analyses revealing downregulated GABAergic signaling, we explored pharmacological strategies to enhance GABA_A_ receptor function in our mouse model. To further advance translational relevance, we also evaluated the pharmacological approach in human brain organoids harboring an *SCN2A* nonsense mutation. Collectively, these experiments aim to clarify the role of Na_V_1.2 in regulating synaptic balance and social behavior and to identify potential therapeutic strategies for *SCN2A*-associated neurodevelopmental disorders.

## RESULTS

### Mice deficient in Na_V_1.2 expression exhibit social impairments

To investigate the behavioral consequences of severe Na_V_1.2 deficiency, we employed a gene-trap (gt) mouse model in which the Na_V_1.2 channel expression is largely reduced^19,20^. Homozygous *Scn2a^gt/gt^* mice (hereafter referred to as HOM or Na_V_1.2-deficient mice) survive to adulthood yet express only ∼25% of the wild-type (WT) Na_V_1.2 level^19^. Because the gene-trap cassette incorporates a *LacZ* reporter driven by the endogenous *Scn2a* promoter (**Figure 1A**)^21,22^, *LacZ*-staining was used as a surrogate marker to delineate the distribution of Na_V_1.2 in the brain. Our results confirm widespread *LacZ* expression, including robust labeling in the striatum (**Figure 1B**), in agreement with previous studies^20,23–25^.

**Figure 1.**
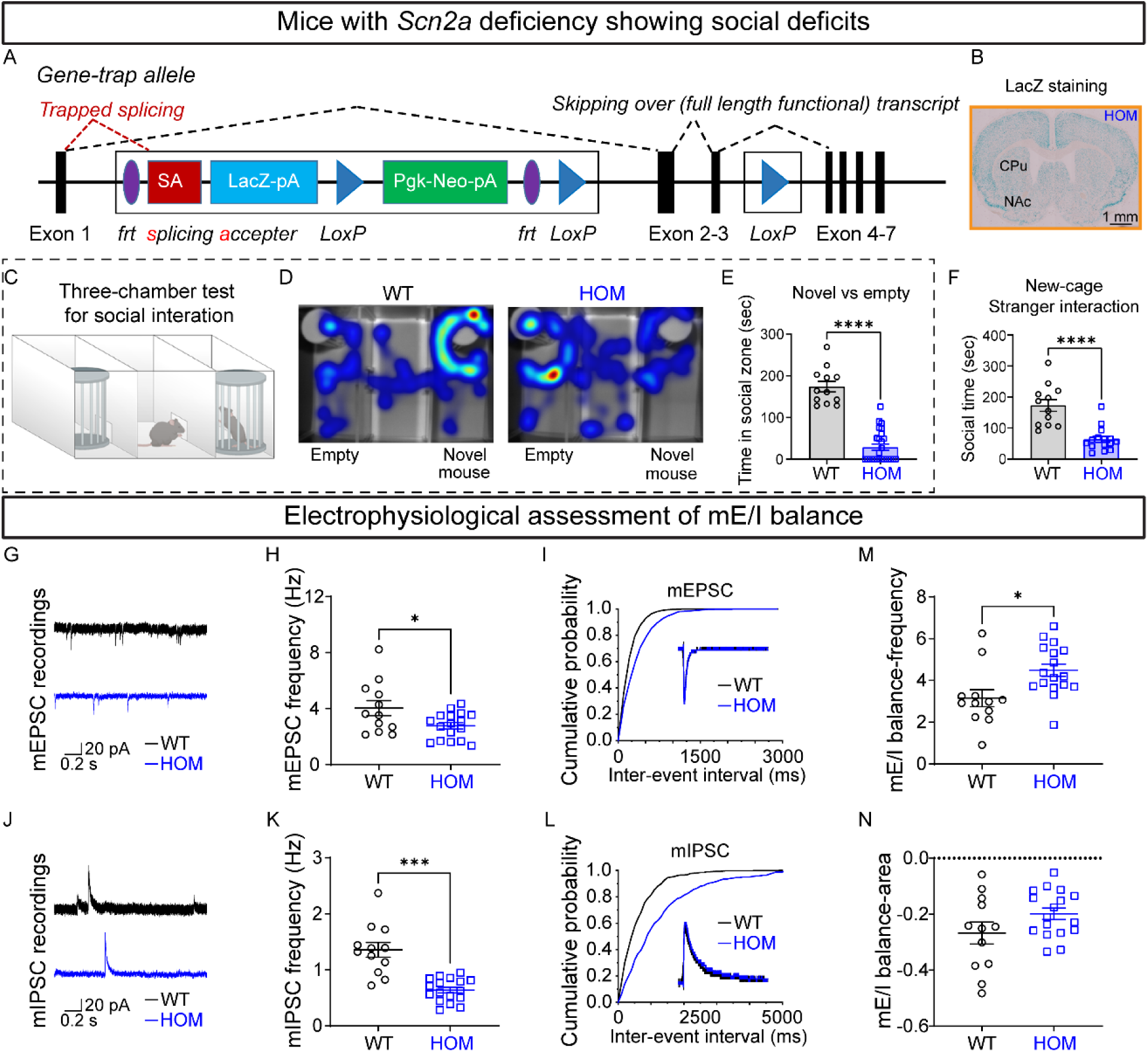
*Scn2a* deficiency results in social deficits and an increased mE/I ratio of medium spiny neurons in the nucleus accumbens. (A) Schematic of the gene trap (gt) allele design. The gt allele has an inserted trapping cassette between the *Exon 1* and *Exon 2* of the *Scn2a* gene in the genome, which traps the transcription from Exon 1 to the trapping cassette, resulting in a deficiency of *Scn2a*. *frt*, *Flp* recognition target (purple); *SA*, splicing acceptor (red); *LacZ*, *lacZ* β-galactosidase (light blue); *LoxP*, locus of X-over P1 (dark blue); and *Neo*, neomycin (green). (B) gt cassette contains a *LacZ* element and is driven by the native *Scn2a* promoter. Thus, the *LacZ* expression can be used as a surrogate of *Scn2a* expression. Representative *LacZ* staining of a coronal slice from a *Scn2a^gt/gt^* (HOM) mouse showing a strong blue signal across the brain including the striatum: caudate putamen (CPu) and nucleus accumbens (NAc). (C) Diagram of a three-chamber social preference test, in which a mouse in the middle chamber can choose to interact with a novel mouse or an empty holding container. (D) Representative heatmaps of tracking trajectories for a single social preference (novel mouse versus empty) trial of *Scn2a^+/+^* (WT) and HOM mice (top view of setup). Warm colors (red, orange, yellow) represent more frequented areas whereas cool colors (green and blue) are less frequented areas. (E) Cumulative time spent in the chamber with the novel mouse during sociability test. n = 12 WT and n = 26 HOM mice. Unpaired t test: ****p < 0.0001. (F) Cumulative social time spent in a clean cage with a novel WT stimulus mouse (stranger). n = 12 WT and n = 15 HOM mice. Unpaired Mann-Whitney *U* test: ****p < 0.0001. (G) Example traces of miniature excitatory postsynaptic currents (mEPSCs) recorded on medium spiny neurons (MSNs) from WT (upper, black) and HOM (lower, blue) mice. (H) mEPSC frequency (n = 12 cells from 6 WT mice and n = 17 cells from 7 HOM mice). Unpaired Welch’s t test: *p < 0.05. (I) Cumulative probability of mEPSC inter-event intervals. Inset: representative averaged mEPSCs. (J) Example traces of miniature inhibitory postsynaptic currents (mIPSCs) recorded on MSNs from WT (upper, black) and HOM (lower, blue) mice. (K) mIPSC frequency (n = 12 cells from 6 WT mice and n = 17 cells from 7 HOM mice). Unpaired Welch’s t test: ***p < 0.001. (L) Cumulative probability of mIPSCs inter-event intervals. Inset: representative averaged mIPSCs. (M) mEPSC/mIPSC ratio calculated with frequency. Unpaired t test: *p < 0.05. (N) mEPSC/sIPSC ratio calculated with area under the curve. Student’s t test: ns, not significant. Data are represented as mean ± SEM.

Social impairment is a central feature of autism spectrum disorder (ASD). Mice typically exhibit high levels of sociability, and social deficits are often evident in mouse models related to autism^26,27^. *Scn2a^+/−^* mice have been reported to display normal or even slightly enhanced sociability relative to WT mice^28,29^. However, our findings reveal that Na_V_1.2-deficient mice display severe social deficits. In the classical three-chamber sociability assay (**Figure 1C**), WT mice spent an average of 174.1 ± 12.8 seconds in the chamber containing a novel stimulus mouse during a 5-minute session, consistent with literature^30^. In contrast, HOM mice spent only 28.7 ± 7.3 seconds in the social chamber (p < 0.0001, unpaired t-test), with no significant sex differences observed (**Figure S1–1A–D**). Heterozygous *Scn2a^+/gt^* (HET) mice did not differ significantly from WT controls in social preference (**Figure S1–1A–D**) like *Scn2a^+/−^* mice^28,29^. In a variant of the three-chamber test, where mice chose between chambers containing a toy mouse and a live novel mouse (**Figure S1–1O**), HOM mice again demonstrated significantly reduced social interaction (WT, 175.4 ± 11.7 s; HET, 151.4 ± 17.1 s, not significant; HOM, 30.62 ± 12.37 s, p < 0.0001, Kruskal–Wallis test) with no apparent sex differences (**Figure S1–1Q)**. WT and HET mice preferred the chamber with the live mouse over both the toy and the neutral middle chamber, whereas HOM mice spent disproportionately more time with the toy mouse, suggesting a pattern of social avoidance (**Figure S1–1P)**.

To further assess the social deficits associated with *Scn2a* deficiency, we conducted several other social behavioral assays. In the reciprocal interaction test, evaluating the latency to initiate social contact (**Figure S1–1L**), number of social bouts (**Figure S1–1M**), and total social time (**Figure S1–1N**) between same-sex pairs of isogenotype (e.g, WT-WT pair or HOM-HOM pair) in a clean cage, HOM mice again exhibited significant impairments (**Figure S1–1L– N**). These deficits were further demonstrated in the stranger interaction test, where interactions between the test mouse (WT or HOM) and a WT stimulus mouse were quantified (**Figures 1F; S1–1E–G**). In the resident-intruder assay, which assesses the aggressive behavior of males by introducing a novel WT mouse into the home cage of a singly-housed resident (WT or HOM), both offensive and defensive behaviors were diminished in HOM mice (**Figure S1–1T–V**). Moreover, continuous monitoring of social interactions in a semi-naturalistic home-cage setting over a 24-hour period revealed significantly reduced social time in HOM mice during both light and dark cycles, irrespective of sex (**Figure S1–1H–K**). Notably, while HET mice generally displayed normal social behaviors, a slight reduction in the number of social bouts was observed, particularly among female HETs during stranger interactions (**Figure S1–1E, G**). Additionally, open-field testing indicated that HOM mice exhibited increased turning behavior and spent more time along the edge of the arena away from the center (**Figure S1–1R, S**), reflecting an anxiety-like phenotype^31^. In summary, our multi-assay behavioral analysis demonstrates that severe Na_V_1.2 deficiency results in robust and multifaceted social impairments. These findings underscore the critical role of Na_V_1.2 channels in the regulation of social behavior and provide a valuable model for investigating the neural mechanisms underlying social deficits.

### Severe *Scn2a* deficiency alters the excitation/inhibition balance in medium spiny neurons

Imbalances in excitatory/inhibitory (E/I) transmission, quantified as the ratio of excitatory postsynaptic currents (EPSCs) to inhibitory postsynaptic currents (IPSCs), have been implicated as a central mechanism underlying autism spectrum disorder (ASD)^15^. Although many ASD mouse models exhibit an elevated E/I ratio^32^, the specific alterations in E/I balance resulting from autism-associated *Scn2a* deficiency remain poorly understood. To assess the impact of severe *Scn2a* deficiency on synaptic transmission, we performed patch-clamp recordings on medium spiny neurons (MSNs) in the nucleus accumbens (NAc) core, a center node in brain circuits critically associated with social behaviors^13,14^. Under pharmacological blockage of inhibitory transmission, the cumulative probability distributions of miniature EPSCs (mEPSCs) in HOM mice shifted toward longer inter-event intervals (**Figure S1–2A–D**), indicating a significant reduction in mEPSC frequency. Surprisingly, a similar reduction with prolonged inter-event intervals was observed for miniature IPSCs (mIPSCs) in HOM MSNs (**Figure S1–2E–H**). When recordings of mEPSCs (**Figure 1G**) and mIPSCs (**Figure 1J**) were obtained from the same neurons, HOM mice exhibited a reduced mEPSC frequency (**Figure 1H, I**) and an even more pronounced decrease in mIPSC frequency (**Figure 1K, L**), resulting in an elevated mE/I ratio compared with WT controls (**Figure 1M**). Because the waveforms of mEPSCs and mIPSCs were unchanged (**Figure 1I, L**), the spike area-based mE/I ratio remained unaltered (**Figure 1N**). Moreover, analysis of spontaneous EPSCs (sEPSCs) and IPSCs (sIPSCs), which include action potential-driven events, also revealed an increased sE/I balance in Na_V_1.2-deficient neurons (**Figure S1–2I–P**). Together, these data indicate that severe *Scn2a* deficiency, while reducing both excitatory and inhibitory inputs, results in a net increase in the basal E/I ratio. This altered balance may disrupt *in vivo* neuronal activity and ultimately contribute to the observed social deficits.

### Enhanced basal *in vivo* neuronal activity and impaired signal-to-noise ratio in Na_V_1.2-deficient striatal neurons

To elucidate how an altered E/I ratio influences basal neuronal activity *in vivo*, we conducted Neuropixels recordings, a cutting-edge high-density electrophysiological technique that allows simultaneous monitoring of hundreds of neurons with high spatial and temporal resolution^33^. The Neuropixels recordings were performed in the striatum of head-fixed mice (**Figure 2A–C**). Raster plots and firing rate analyses demonstrated that HOM mice exhibit significantly enhanced basal firing compared with WT controls (**Figure 2D, E**). To understand how *Scn2a* deficiency differentially affects distinct neuronal populations, neurons were classified using electrophysiological criteria, including waveform duration, long interspike intervals, and post-spike suppression, allowing segregation into putative MSNs, fast-spiking interneurons (FSIs), tonically active neurons (TANs), and unidentified interneurons (UINs)^34^ (**Figures 2F; S2A(i, ii)–B**). Notably, the firing rates of MSNs, the principal striatal neurons, were markedly elevated in HOM mice (**Figures 2G, H; S2C**). FSIs showed a moderate increase in firing rate (**Figure S2D**), and TANs as well as UINs were unaffected (**Figure S2E, F**), indicating that *Scn2a* deficiency predominantly drives hyperactivity in striatal principal neurons. Given that MSNs do not spontaneously fire in brain slices^20^, this *in vivo* hyperactivity is likely attributable to an elevated E/I ratio. Because the neuronal firings were recorded under conditions unrelated to social behavior, we considered this elevated basal activity to represent increased “noisy” neural signaling.

**Figure 2.**
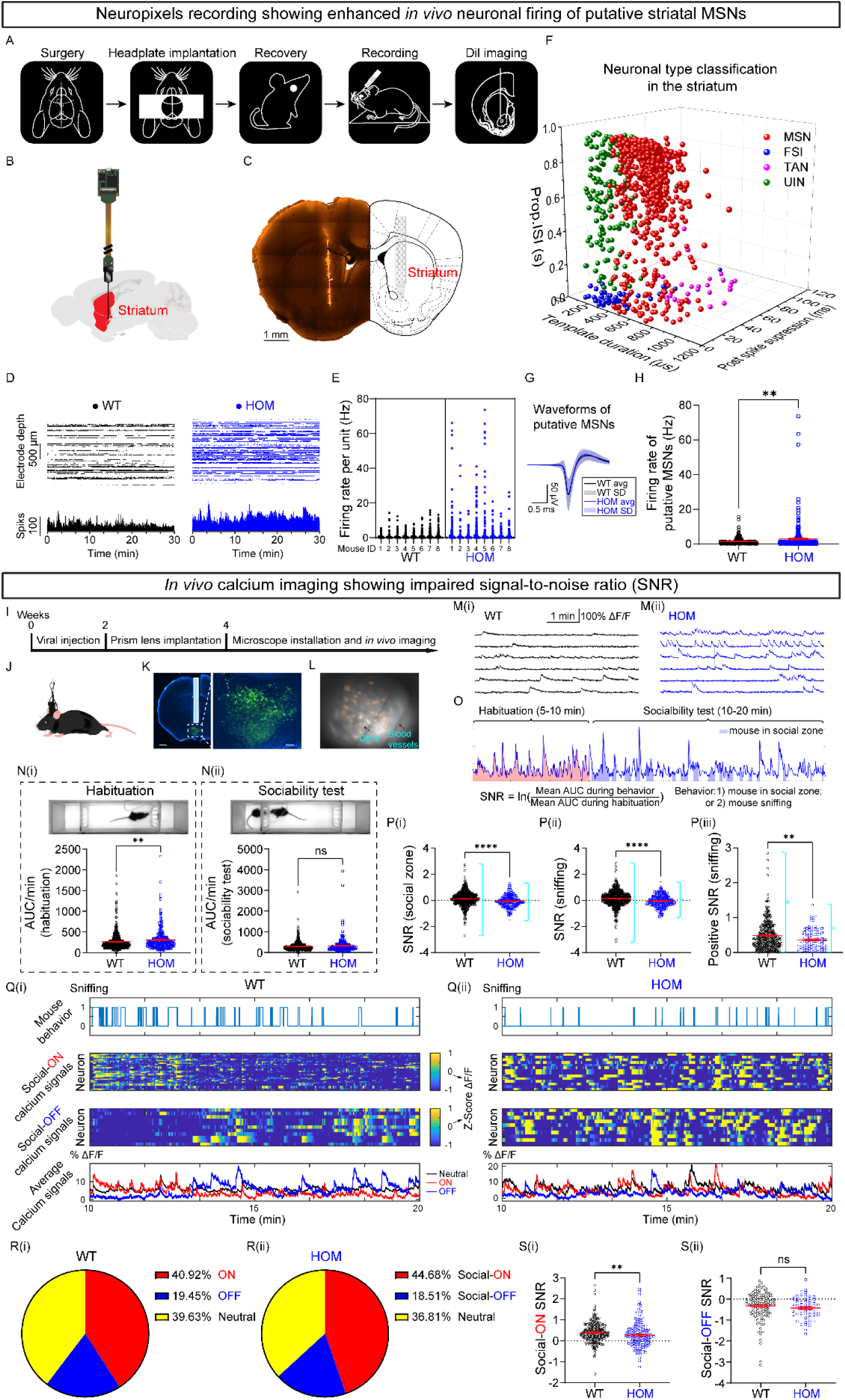
Enhanced basal *in vivo* neuronal activity is linked to impaired SNR due to *Scn2a* deficiency in mice. (A–H) Neuropixels recordings of neuronal activity in the striatum. (A) Experimental timeline, including surgery, headplate implantation, recovery, Neuropixels recording, and DiI imaging for post recording validation. (B) Schematic of the Neuropixels probe insertion into the striatum (red region). (C) Coronal section showing Neuropixels probe placement in the striatum with the Dil imaging post recording. (D) Representative raster plots (top) and corresponding firing rate histograms (bottom; downsampled by a factor of 10) recorded from WT (black) and *Scn2a^gt/gt^* (HOM, blue) mice. (E) Quantification of firing rates per unit across individual WT and HOM mice. (F) Neuronal classification within the striatum based on electrophysiological properties. Striatal cells were identified as putative medium spiny neurons (MSNs), fast-spiking interneurons (FSIs), tonically active neurons (TANs), and a fourth class of unidentified interneurons (UINs), according to waveform duration, length of post-spike suppression, and fraction of long interspike intervals. (G) Average waveforms of putative MSNs recorded in WT and HOM mice. (H) Firing rates of putative MSNs. n = 288 MSNs from 8 WT mice and n = 314 MSNs from 8 HOM mice. Linear mixed-effects model fit by maximum likelihood: **p < 0.01. Data are presented as mean ± SEM. (I–S) *In vivo* calcium imaging of neuronal activity during social behaviors. (I) Experimental timeline, including viral injection, prism lens implantation, microscope installation, and *in vivo* imaging. (J) Representative setup for calcium imaging in freely moving mice. (K) Representative coronal brain slice showing viral expression of the genetically encoded calcium indicator GCaMP6s in striatal neurons (green) and location of GRIN lens track (light blue). Scale bar = 500 μm. Zoomed-in view of the region outlined by the white dashed rectangle is shown on the right. Scale bar = 100 μm. Nuclei are stained with DAPI (blue). (L) Example field of view. A clear blood vessel pattern is visible with some cells in the image. (M) Representative extracted ΔF/F Ca^2+^ transients from WT (M(i)) and HOM (M(ii)) mice during habituation. Horizontal scale bar 1 minute and vertical scale bar 100% ΔF/F. (N) Quantification of neuronal activity, measured as area under the curve (AUC) during habituation (N(i)) and sociability tests (N(ii)). HOM mice exhibit significantly increased neuronal activity compared with WT during habituation, but not during sociability test. n = 623 cells from 3 WT mice and n = 307 cells from 5 HOM mice. Unpaired Mann-Whitney *U* test: ns, not significant; **p < 0.01. (O) Schematic showing SNR analysis algorithm. (P) SNR quantification when mouse in social zones (P(i)) and during sniffing events only (P(ii) and P(iii)), showing significantly reduced SNR in HOM mice. n = 626 (390 positive sniffing) cells from 3 WT mice and n = 228 (103 positive sniffing) cells from 3 HOM mice. Unpaired Mann-Whitney *U* test: **p < 0.01; ****p < 0.0001. (Q) Example social-ON and social-OFF neurons, identified based on the similarity between sniffing behavior and Ca^2+^ activity, recorded in the striatum from WT (Q(i)) and HOM (Q(ii)) mice. Top panel: blue lines indicate sniffing behavior epochs. Middle panels: heatmaps show normalized Ca^2+^ traces of individual social-ON and social-OFF neurons. Bottom panel: averaged Ca^2+^ traces of ON, OFF, and Neutral neurons shown separately. (R) Pie charts showing the distribution of social-ON, social-OFF, and neutral neurons from WT (R(i)) and HOM (R(ii)) mice. (S) Quantification of social-ON (S(i)) and social-OFF (S(ii)) SNR, revealing a significant impairment in SNR for social-ON neurons (shifted towards smaller positive values) in HOM mice, but not for social-OFF neurons, which do not exhibit a shift towards less negative values. n = 260 social-ON and 123 social-OFF cells from 3 WT mice; and n = 147 social-ON and 58 social-OFF cells from 3 HOM mice. Unpaired Mann-Whitney *U* test: ns, not significant; **p < 0.01. Data are represented as mean ± SEM.

We next examined how increased basal “noisy” neuronal activity affects the encoding of behaviorally relevant neuronal signals during social interactions, by performing *in vivo* calcium imaging during social behaviors, as Neuropixels recordings in freely moving mice remain technically challenging. Viral delivery of GCaMP6s into the striatum, combined with GRIN lens implantation (**Figure 2I–K**), allowed us to monitor neuronal Ca^2+^ transients in freely moving mice (**Figure 2L**). During the habituation period, ΔF/F traces from HOM mice revealed enhanced activity profiles, characterized by larger calcium transient amplitudes and faster risetimes relative to WT, while decay times remained comparable (**Figures 2M; S2G–J**). This observation was further supported by area-under-the-curve (AUC) analyses, which showed a significant increase in HOM mice during habituation (**Figure 2N(i)**), consistent with the hyperexcitability observed via Neuropixels recordings (**Figure 2A–H**). However, during sociability test, differences in AUC largely disappeared (**Figure 2N(ii)**), and only faster risetimes and larger amplitudes persisted (**Figure S2K–N**). This pattern suggests the *Scn2a*-deficient neurons are unable to effectively adjust their responses from the elevated basal “noisy” activity to social interactions.

To quantify the fidelity of behavioral signal encoding, we computed the signal-to-noise ratio (SNR), as schematically represented in **Figure 2O**. Specifically, for each neuron, we defined “noise” as neuronal activity during habituation, and “signal” as activity in response to social stimuli. When mice occupied social zones, particularly during active sniffing events, the SNR was significantly reduced in HOM mice relative to WT (**Figures 2P(i–iii); S2O–P**). Moreover, the narrow range of distribution of neuronal activity in SNR values observed in HOM mice suggests a diminished responsiveness to social stimuli. For clarity, we defined social-ON neurons as those that exhibit increased Ca^2+^ activity in response to social interactions, whereas social-OFF neurons display a decrease in activity under the same conditions. Further analysis of these subpopulations during sniffing revealed that altered Ca^2+^ dynamics likely underlie impaired social signal discrimination in HOM mice (**Figure 2Q–S**). Collectively, these findings indicate that *Scn2a* deficiency not only elevates basal “noisy” activity but also impairs the neurons’ ability to generate robust, behaviorally relevant signals.

### Global restoration of *Scn2a* in adulthood rescues E/I imbalance and social behavior

To determine whether restoration of *Scn2a* expression in adulthood can normalize synaptic transmission and rescue social deficits, we employed a genetic rescue strategy by Flp-mediated excision of the trapping cassette from the *gt* allele (**Figure 3A**). To achieve global transduction, we utilized AAV-PHP.eB, a capsid variant developed with enhanced ability to cross the blood-brain barrier. Adult WT and HOM mice received tail-vein injections of AAV-PHP.eB encoding either optimized Flp recombinase (Flp) or a control vector (**Figure 3J**). In HOM mice treated with AAV-Flp, deletion efficiency of the trapping cassette was confirmed by *LacZ* staining in the CPu and NAc (**Figure 3K**), and Western blot analysis demonstrated partial restoration of Na_V_1.2 expression (**Figure 3L**)^35^.

**Figure 3.**
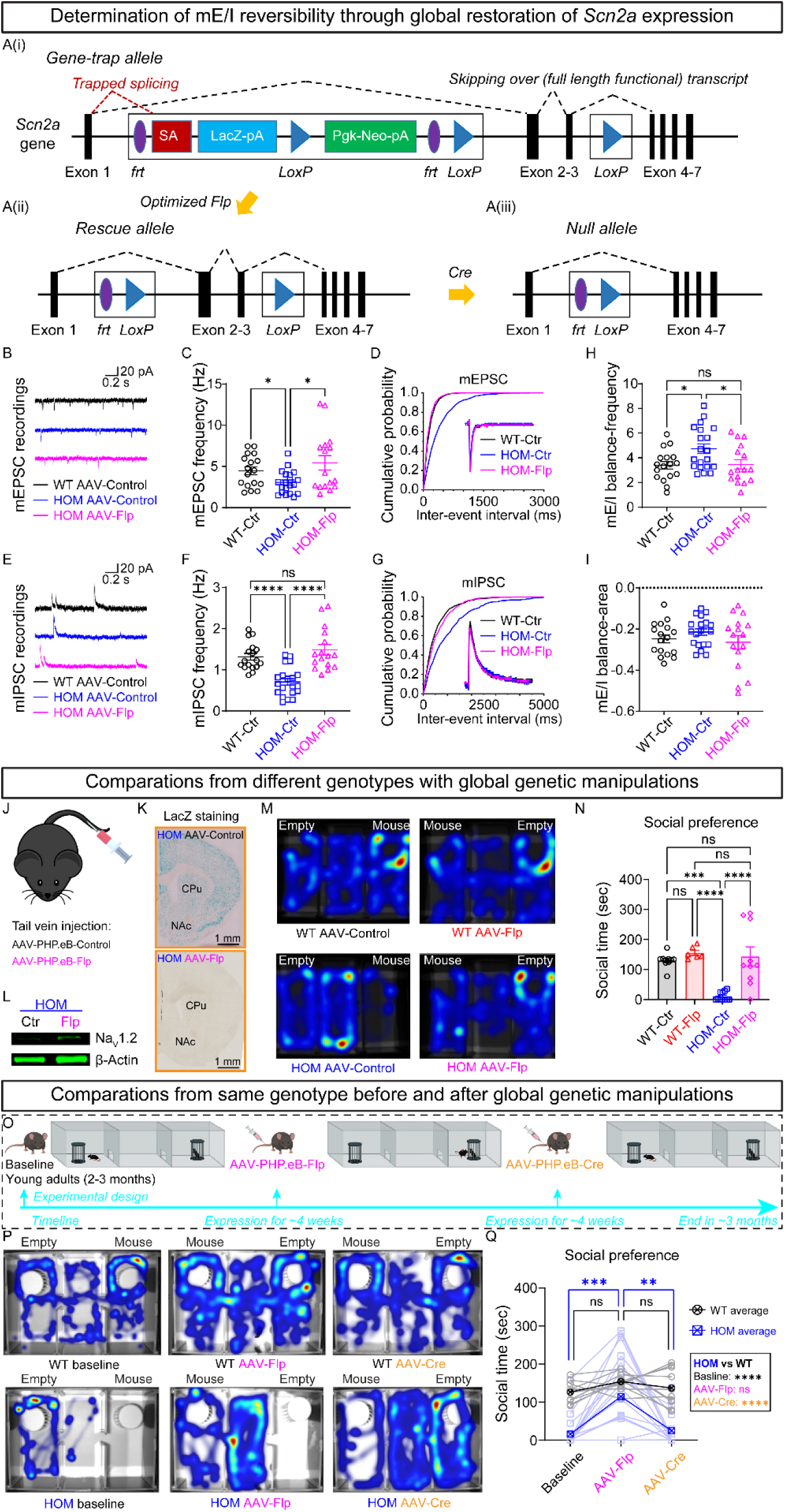
Global restoration of *Scn2a* expression in adulthood reverses mE/I ratio and rescues social impairment in Na_V_1.2-deficient mice. (A) Schematic design of the *gt* allele (Ai), rescue allele following Flp recombination (Aii), and null allele after Cre-mediated recombination (Aiii). In the presence of *Flp* recombinase, *frt* sites flanked trapping cassette will be removed, producing a rescue allele that allows the expression of *Scn2a* towards the WT level. In the presence of *Cre* recombinase, *LoxP* sites flanked *Exon* 2-3 will be removed, producing a complete knockout (null) allele. (B–D) Analysis of mEPSCs recorded on MSNs from WT mice transduced with AAV-Control (n = 17 cells from 6 mice, black), HOM mice transduced with AAV-Control (n = 20 cells from 7 mice, blue), and HOM mice transduced with AAV-Flp (n = 16 cells from 6 mice, magenta). (B) Representative mEPSC traces, (C) frequency, and (D) cumulative probability of inter-event intervals (Inset: representative averaged mEPSCs). Brown-Forsythe and Welch one-way ANOVA with Dunnett T3’s multiple comparisons: *p < 0.05. (E–G) Analysis of mIPSCs recorded on MSNs from WT mice transduced with AAV-Control (n = 17 cells from 6 mice, black), HOM mice transduced with AAV-Control (n = 20 cells from 7 mice, blue), and HOM mice transduced with AAV-Flp (n = 16 cells from 6 mice, magenta). (E) Representative mIPSC traces, (F) frequency, and (G) cumulative probability of inter-event intervals (Inset: representative averaged mIPSCs). Brown-Forsythe and Welch one-way ANOVA with Dunnett T3’s multiple comparisons: ns, not significant; ****p < 0.0001. (H, I) Ratios of mE/I by frequency (H) and area (I) highlighting restored E/I balance in HOM mice transduced with AAV-Flp. One-way ANOVA with Tukey’s multiple comparisons (M, N): ns, not significant; *p < 0.05. (J) Schematic tail vein injection of AAV-PHP.eB-Flp or AAV-PHP.eB-Control in WT and *Scn2a^gt/gt^* (HOM) mice. (K) *LacZ* staining in the CPu and NAc regions confirms trapping cassette deletion in HOM mice injected with AAV-PHP.eB-Flp. (L) Western blot analysis of Na_V_1.2 and β-actin levels in WT and HOM mice. (M) Representative heatmaps of tracking trajectories for a single social preference (novel mouse versus empty) trial of WT and HOM mice injected with AAV-Control or AAV-Flp (top view of set up). Warm colors (red, orange, yellow) represent more frequented areas whereas cool colors (green and blue) are less frequented areas. (N) Cumulative time spent in the chamber with the novel mouse during sociability test. n = 10 WT mice transduced with AAV-Control, n = 6 WT mice transduced with AAV-Flp, n = 11 HOM mice transduced with AAV-Control, and n = 10 HOM mice transduced with AAV-Flp. One-way ANOVA with Bonferroni’s multiple comparisons: ns, not significant; ***p < 0.001; ****p < 0.0001. (O) Experimental timeline for assessing social preference, before and after AAV-Flp and AAV-Cre transduction. (P) Heatmaps of baseline and post-injection social interactions from WT and HOM mice treated with AAV-Flp or AAV-Cre. (Q) Quantification of social preference before and after AAV-Flp and AAV-Cre sequential transduction. n = 10 WT and n = 19 HOM mice. Inset: comparisons between genotypes in the same treatment condition. Two-way ANOVA with Tukey’s multiple comparisons: ns, not significant; **p < 0.01; ***p < 0.001; ****p < 0.0001. Data are represented as mean ± SEM.

To investigate the synaptic basis of the rescue, we performed whole-cell recordings on striatal MSNs. In slices from HOM mice transduced with AAV-Flp, the reduced mEPSC and sEPSC frequencies were normalized to levels comparable to WT (**Figures 3B–D; S3–1A–C**). Similarly, the decreased inhibitory transmissions observed in HOM mice were restored following Flp treatment, as indicated by analyses of mIPSCs and sIPSCs (**Figures 3E–G; S3–1D–F**). Consequently, the E/I ratio, particularly assessed by event frequency, was normalized towards WT levels in HOM mice after AAV-Flp injection (**Figures 3H; S3–1G, H**).

Behaviorally, HOM mice injected with AAV-Control displayed markedly reduced exploration of the chamber containing a novel mouse in the three-chamber sociability assay (**Figure 3M, N**). In contrast, HOM mice receiving AAV-Flp spent significantly more time in the novel mouse chamber, reaching levels comparable to WT controls (**Figure 3M, N**; WT-Control 130.1 ± 7.3 s, WT-Flp 155.4 ± 8.3 s, HOM-Control 10.5 ± 4.3 s, HOM-Flp 143.1 ± 32.0 s; p < 0.001, p < 0.0001, one-way ANOVA). As expected, Flp-treated WT mice exhibited similar social behaviors to those of control-treated WT animals (**Figure 3M, N**). Notably, a lower dose of Flp alleviated social impairments primarily in females, leaving male HOM mice unrescued in the three-chamber test (**Figure S3–2A–C**), which indicates a dose-related effect with females being more responsive. Importantly, the restoration of Na_V_1.2 selectively improved social behavior only, while other parameters, such as open field distance, anxiety-like behavior, marble burying, and nesting, remained unaffected with treatment in adulthood (**Figures S3–2D–I**; **S3–1I–N**). Together, these results indicate that the global restoration of Na_V_1.2 in adulthood may be sufficient to rescue sociability.

Finally, a sequential transduction experiment (**Figure 3O**) further validated the robustness of the rescue. Baseline heatmaps confirmed social interaction deficits in HOM mice, which were ameliorated following Flp-mediated restoration of *Scn2a* expression (**Figure 3P**). Quantitative analysis revealed that while untreated HOM mice exhibited significant social impairments relative to WT, global restoration of *Scn2a* fully rescued social behavior (**Figure 3Q**). Remarkably, the subsequent depletion of *Scn2a* (null) in adulthood, after the global restoration of *Scn2a* following the injection of Cre recombinase (**Figure 3A(i–iii)**), recapitulated the social deficits, reinforcing the necessity of *Scn2a* expression for maintaining normal social behaviors (**Figure 3Q**). Together, these data demonstrate that restoring *Scn2a* expression in adulthood reverses the imbalanced excitatory-inhibitory synaptic transmission in the striatum and rescues social deficits in Na_V_1.2-deficient mice.

### Selective restoration of *Scn2a* in NAc-projecting neurons lowers basal noisy activity and rescues social behavior

To move beyond a global rescue and pinpoint the key brain region mediating the rescue effect, we examined whether targeting the striatum, a common node mediating social behaviors^13,14^, could ameliorate aberrant basal activity and social deficits. Notably, we previously found that *Scn2a* deficiency induces unexpected intrinsic hyperexcitability in striatal MSNs^20^. To determine whether this intrinsic hyperexcitability or the E/I imbalance drives social impairments, we selectively restored Na_V_1.2 either in the striatum or its afferents. We employed two viral strategies (**Figure 4A**): direct transduction of striatal neurons using AAV-PHP.eB-Flp (**Figure 4B, top**) and retrograde transduction of striatum-projecting neurons using AAVrg-Flp (**Figure 4B, bottom**). Viral imaging confirmed the robust targeting of projection neurons in the mPFC, S1, and insular cortex (**Figure 4B, bottom**).

**Figure 4.**
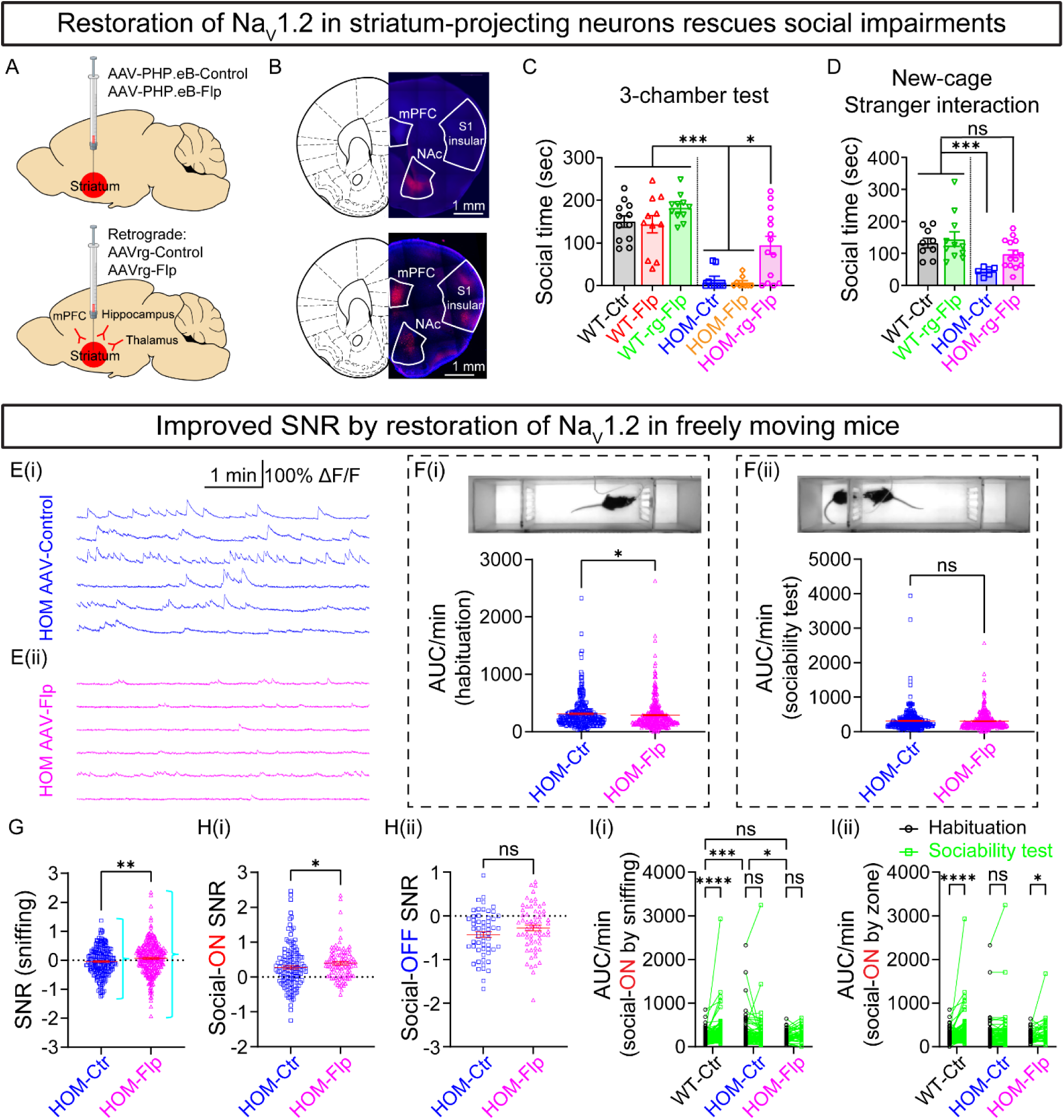
Lowered noisy basal neuronal activity by restoration of *Scn2a* expression in NAc-projecting neurons improves SNR and rescues social deficits in adult Na_V_1.2-deficient mice. (A) Schematic of targeted viral delivery to striatal neurons (top) and striatum-projecting neurons (bottom) using AAV-PHP.eB-Flp or AAVrg-Flp (retrograde), respectively. (B) *LacZ* staining confirms viral targeting in striatum-projecting neurons located in upstream regions (e.g. mPFC, S1, and insular cortex). Scale bar = 1 mm. (C) Three-chamber sociability test showing social time spent in the chamber with the novel mouse (novel mouse versus empty). n = 12 WT mice transduced with AAV-Control (black), n = 11 WT mice transduced with AAV-Flp (red), n = 11 WT mice transduced with retrograde AAV-Flp (green), n = 9 HOM mice transduced with AAV-Control (blue), n = 9 HOM mice transduced with AAV-Flp (orange), and n = 14 HOM mice transduced with retrograde AAV-Flp (magenta). One-way ANOVA with Bonferroni’s multiple comparisons: *p < 0.05; ***p < 0.001. (D) Cumulative social time spent in a clean cage with a novel WT stimulus mouse (stranger). n = 9 WT mice transduced with AAV-Control (black), n = 11 WT mice transduced with retrograde AAV-Flp (green), n = 6 HOM mice transduced with AAV-Control (blue), and n = 14 HOM mice transduced with retrograde AAV-Flp (magenta). One-way ANOVA with Bonferroni’s multiple comparisons: ns, not significant; ***p < 0.001. (E) Representative extracted ΔF/F Ca^2+^ transients from HOM mice transduced with AAV-Control ((Ei), blue), and HOM mice transduced with AAV-Flp ((Eii), magenta) during habituation. Horizontal scale bar 1 minute and vertical scale bar 100% ΔF/F. (F) Quantification of neuronal activity, measured as area under the curve (AUC) during habituation (F(i)) and sociability tests (F(ii)). HOM-Flp mice exhibit decreased activity compared with HOM-Ctr during habituation, but not during sociability test. n = 307 cells from 5 HOM mice transduced with AAV-Control and n = 360 cells from 8 HOM mice transduced with retrograde AAV-Flp. Unpaired Mann-Whitney *U* test: ns, not significant; *p < 0.05. (G) Quantification of sniffing-related SNR, demonstrating elevated SNR in HOM-Flp mice. n = 228 cells from 3 HOM mice transduced with AAV-Control and n = 296 cells from 6 HOM mice transduced with retrograde AAV-Flp. Unpaired Welch’s t test: **p < 0.01. (H) Quantification of social-ON (H(i)) and social-OFF (H(ii)) SNR, showing an improvement in SNR for social-ON neurons (shifted towards larger positive values) in HOM-Flp mice, but not for social-OFF neurons, which do not exhibit a shift towards more negative values. n = 147 social-ON and 58 social-OFF cells from 3 HOM mice transduced with AAV-Control; and n = 101 social-ON and 65 social-OFF cells from 6 HOM mice transduced with retrograde AAV-Flp. Unpaired Mann-Whitney *U* test (H(i)) and unpaired t test (H(ii)): ns, not significant; *p < 0.05. (I) Quantification of neuronal activity of social-ON neurons, analyzed by sniffing (I(i)) and by zone-based social behavior (I(ii)), during habituation (black) and sociability tests (green). HOM-Flp mice exhibited a normalization of the enhanced activity observed in HOM-Ctr mice during habituation, and showed improved neuronal responsiveness during sociability sociability test. Sniffing: n = 207 cells from 3 WT mice transduced with AAV-Control, n = 86 cells from 3 HOM mice transduced with AAV-Control, and n = 51 cells from 3 HOM mice transduced with retrograde AAV-Flp. Social zone: n = 197 cells from 3 WT mice transduced with AAV-Control, n = 93 cells from 3 HOM mice transduced with AAV-Control, and n = 47 cells from 3 HOM mice transduced with retrograde AAV-Flp. Paired two-way ANOVA with Tukey’s multiple comparisons: ns, not significant; *p < 0.05; ***p < 0.001; ****p < 0.0001. Data are represented as mean ± SEM.

In the three-chamber sociability assay, WT mice injected with AAV-Control, AAV-Flp, or retrograde AAV-Flp exhibited comparable social exploration times (**Figure 4C**; WT-Control 150.2 ± 13.3 s, WT-Flp 143.6 ± 20.4 s, WT-retrograde-Flp 182.2 ± 9.5 s; not significant by one-way ANOVA). Expectedly, HOM mice injected with AAV-Control showed markedly reduced social interaction. Surprisingly, direct restoration of Na_V_1.2 in striatal neurons via AAV-Flp failed to improve social behavior, suggesting that targeting striatal neurons alone is insufficient, a conclusion supported by complementary experiments using chemogenetic modulations to directly increase or decrease striatal neuron activity. (**Figure S4–1C, D**). In contrast, retrograde delivery of Flp, which targets NAc-projecting neurons, robustly increased the time HOM mice spent in the novel mouse chamber (**Figure 4C**; HOM-Control 13.8 ± 8.3 s, HOM-Flp 7.3 ± 4.3 s, HOM-retrograde-Flp 94.6 ± 21.3 s; p < 0.05, p < 0.001, one-way ANOVA), an effect observed in both sexes (**Figure S4–1A**). The cumulative social time during stranger interactions was also significantly enhanced in HOM mice receiving retrograde AAV-Flp (**Figures 4D; S4–1B**), further suggesting rescue of social behavior. Consistent with global restoration, retrograde AAV-Flp selectively rescued social behaviors without affecting open field turning, center duration, distance, or anxiety-like behaviors in adulthood (**Figure S4–1E–H**).

*In vivo* calcium imaging in HOM mice further elucidated the neuronal activity correlations of this behavior rescue. Representative ΔF/F Ca^2+^ traces during habituation revealed that basal noisy activity was markedly reduced following retrograde AAV-Flp transduction compared with AAV-Control (**Figures 4E, F(i); S4–1I–L**), while AUC and other Ca^2+^ transient properties during sociability tests remained unchanged (**Figures 4F(ii); S4–1M–P**). Analysis of behaviorally relevant neuronal activity demonstrated that sniffing-related SNR was significantly increased in HOM mice treated with retrograde AAV-Flp, accompanied by an expanded SNR distribution range (**Figures 4G; S4–1Q**), indicating an enhancement in neuronal responsiveness to social stimuli. While social-OFF neurons exhibited no significant changes in SNR (**Figures 4H(ii); S4– 1R**), social-ON neurons displayed a marked improvement (**Figure 4H(i)**). To further validate the hypothesis that elevated basal neuronal noise weakens social signaling, we analyzed social-ON neuron activity. HOM mice exhibited significantly higher basal activity, which diminished the neuronal response to social stimuli observed in WT mice (**Figure 4I(i)**). However, retrograde AAV-Flp transduction in HOM mice reversed this increased basal activity (**Figure 4I(i)**) and restored responsiveness to social stimuli to WT-like levels (**Figure 4I(ii)**). Together, these results indicate that selective restoration of *Scn2a* in NAc-projecting neurons reduces basal noisy Ca^2+^ activity and normalizes the SNR during social interactions, thereby rescuing social deficits.

To investigate how *Scn2a* expression influences neuronal activity and behavioral coding, we further bidirectionally modulated its expression in these viral-injected mice via subsequent systemic AAV transduction. In HOM-Ctr mice (control AAV into striatum), *Scn2a* restoration through tail-vein AAV-Flp injection significantly increased calcium transient AUC, rate, and amplitude during the sociability test (**Figure S4–2A–E**). This treatment also improved SNR, selectively enhancing social-ON neuronal signaling while leaving social-OFF neurons unchanged (**Figure S4–2F–G**). Conversely, *Scn2a* depletion via tail-vein injection of AAV-Cre in HOM-Flp mice (retrograde Flp into striatum) altered transient dynamics (**Figure S4–2H–L**) and induced an upshifted negative SNR component along with impaired social-OFF neuron signaling during sniffing events (**Figure S4–2M–N**). These findings suggest that *Scn2a* restoring discriminatively enhances social-ON signal processing by reducing basal noise, whereas its further suppression exacerbates aberrant neuronal activity and disrupts social-OFF signal encoding.

### GABAergic dysregulation and pharmacological enhancement of inhibitory transmission reduces basal noisy activity in both mouse and human models of *SCN2A* deficiency

Given the predominant reduction in IPSCs observed in Na_V_1.2-deficient mice (**Figures 1G–N; S1–2**), we uncovered the molecular mechanisms underlying *Scn2a* deficiency-associated phenotypes through RNA sequencing. Focusing on GABA pathway-related genes, KEGG pathway analysis revealed a pronounced downregulation of the GABA signaling pathway in mice with severe *Scn2a* deficiency (**Figure S5A**)^36^. Furthermore, a volcano plot comparing HOM with WT brain samples identified numerous significantly decreased transcripts of GABA pathway-related genes (**Figure 5A**), a result corroborated by the corresponding heatmap (**Figure 5B**).

**Figure 5.**
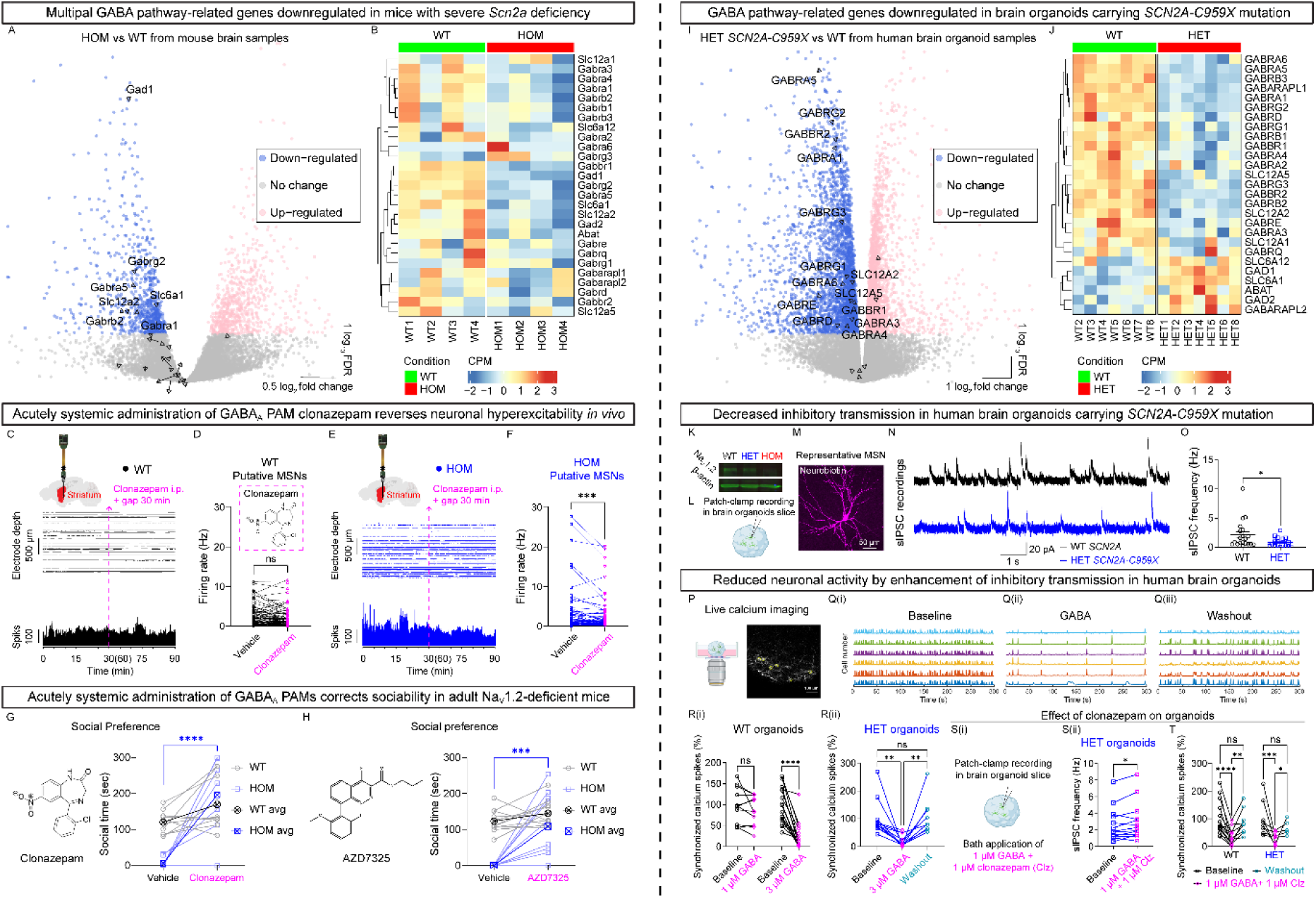
GABAergic dysregulation and pharmacological enhancing inhibitory transmission lowers basal nosy activity in both mouse and human models with *SCN2A* deficiency. (A, B) RNA sequencing analysis of GABA pathway-related gene expression in mice with severe *Scn2a* deficiency. (A) Volcano plot showing differentially expressed genes in *Scn2a^gt/gt^* (HOM) versus *Scn2a^+/+^* (WT) mouse brain samples. Arrows highlight key GABA pathway-related genes, with significantly downregulated genes annotated by name, whereas few other genes remain unannotated. (B) Heatmap illustrating expression levels of key GABA-related genes across WT and HOM samples. (C–F) Acute systemic administration of clonazepam, a GABA_A_ receptor-positive allosteric modulator (PAM), reverses neuronal hyperexcitability *in vivo*. (C) Schematic depicting Neuropixels probe insertion into the striatum (red region, top), a representative raster plot (middle), and corresponding firing rate histogram (bottom; downsampled by a factor of 10) recorded from WT mice (black). (D) Quantification of firing rate of putative MSNs before and after clonazepam administration in WT mice (n= 96 cells from 3 mice). Inset: schematic of clonazepam chemical structure. (E) Schematic depicting Neuropixels probe insertion into the striatum (red region, top), a representative raster plot (middle) and corresponding firing rate histogram (bottom; downsampled by a factor of 10) of neurons recorded from HOM mice (blue). (F) Quantification of firing rate of putative MSNs before and after clonazepam administration in HOM mice (n= 105 cells from 3 mice). Paired Wilcoxon test: ns, not significant; ***p < 0.001. (G, H) Acute systemic administration of PAMs (i.p., 30 minutes before behavioral test) corrects sociability deficits in adult Na_V_1.2-deficient mice. Quantification of social interaction behaviors in the three-chamber test (novel mouse versus empty) after administration of clonazepam (G, n = 10 mice for each genotype) or AZD7325 (H, n = 11 for WT and n = 10 for HOM mice). Paired two-way ANOVA with Bonferroni’s multiple comparisons: ***p < 0.001; ****p < 0.0001. (I, J) RNA sequencing analysis of GABA pathway-related gene expression in human brain organoids carrying the *SCN2A-C959X* mutation, identified in children with profound autism. (I) Volcano plot showing differentially expressed genes in heterozygous (HET) *SCN2A^+/C959X^* versus WT human brain organoids. Arrows highlight key GABA pathway-related genes, with significantly downregulated genes annotated by name, whereas few other genes remain unannotated. (J) Heatmap illustrating expression levels of key GABA pathway-related genes across WT and HET organoid samples. (K–O) Decreased inhibitory transmission in human brain organoids carrying the *SCN2A-C959X* mutation. (K) Western blot analysis of Na_V_1.2 and β-actin levels in WT, *SCN2A^+/C959X^* (HET), and *SCN2A^C959X/C959X^* (HOM) organoids. (L) Schematic of patch-clamp recordings performed in human brain organoid slices. (M) Representative MSN labeled with neurobiotin-AlexaFluor 647 after patch-clamp recording. Scale bar = 50 μm. (N) Representative sIPSC recordings from WT and HET organoids. Horizontal scale bar 1 second and vertical scale bar 20 pA. (O) Quantification of sIPSC frequency. n = 20 cells from 6 WT organoids and n = 17 cells from 5 HET organoids. Unpaired Mann-Whitney *U* test: *p < 0.05. (P) Schematic of experimental setup for calcium imaging and pharmacological intervention in brain organoids. (Q(i–iii)) Representative traces showing neuronal calcium activity under baseline, GABA administration, and washout conditions. (R(i, ii)) Quantification of normalized synchronized calcium transient rates in WT (n = 11 cells from 4 organoids for 1 μM GABA group; n = 17 cells from 6 organoids for 3 μM GABA group) and HET (n = 12 cells from 4 organoids for 3 μM GABA group). Two-way ANOVA with uncorrected Fisher’s LSD’s multiple comparisons (R(i)), and mixed-effects analysis with Tukey’s multiple comparisons (R(ii)): ns, not significant; **p < 0.01; ****p < 0.0001. (S(i)) Schematic of patch-clamp recordings in brain organoid slices, and (S(ii)) effect of co-application of GABA (1 µM) and clonazepam (Clz, 1 µM) on sIPSC frequency in HET brain organoids. n = 13 cells from 4 HET organoids. Paired t test: **p < 0.01. (T) Normalized synchronized calcium spike rates in WT (n = 14 cells from 6 organoids) and HET (n = 8 cells from 3 organoids) organoids under baseline, Clz, and washout conditions. Mixed-effects analysis with uncorrected Fisher’s LSD: ns, not significant; *p < 0.05; **p < 0.01; ***p < 0.001; ****p < 0.0001. Data are presented as mean ± SEM.

Because reduced GABAergic signaling can enhance the E/I ratio and contribute to *in vivo* neuronal hyperexcitability, we examined whether potentiating GABA_A_ receptor activity using a positive allosteric modulator (PAM) could normalize neuronal firing. Notably, clonazepam, an FDA-approved benzodiazepine has been shown to rescue social deficits in both a *Scn1a* knockout model of Dravet syndrome^37^ and the BTBR model of idiopathic autism^38^. In WT mice, baseline recordings from putative MSNs showed low firing rates that remained unchanged following acute systemic administration of a low dose of clonazepam (Clz, 0.05 mg/kg, i.p.) (**Figure 5C, D**). In contrast, clonazepam markedly suppressed the abnormally high firing rates in HOM mice (**Figure 5E, F**).

Next, to assess whether enhancing GABAergic transmission could rescue social behavioral deficits in adult Na_V_1.2-deficient mice, we tested an additional PAM, AZD7325, currently in clinical trials for anxiety disorders and previously shown to be beneficial in a Dravet model^39^. Acute intraperitoneal administration of clonazepam (0.05 mg/kg, i.p.) or AZD7325 (3 mg/kg, i.p.) significantly improved social interactions in HOM mice in the 3-chamber test (**Figure 5G, H**). Importantly, neither compound induced notable sedative effects in WT or HOM mice, as assessed by the open field test (**Figure S5C–H**). These data suggest that targeting the GABA pathway is sufficient to rescue social deficits in adult Na_V_1.2-deficient mice.

To evaluate the translational potential of these findings, we examined human brain organoids carrying the heterozygous (HET) nonsense mutation *SCN2A* c.2877C>A (p.Cys959Ter), referred to as *SCN2A-C959X*, identified in children with profound ASD^4,40,41^. Western blot analysis confirmed reduced Na_V_1.2 protein levels in HET organoids compared with WT (**Figure 5K**). Consistent with the mouse data, RNA sequencing analysis of *SCN2A-C959X* organoids revealed significant downregulation of GABA pathway-related genes (volcano plot in **Figure 5I**, heatmap in **Figure 5J**, and enriched KEGG pathways in **Figure S5B**). Functionally, patch-clamp recordings in human striatal organoid slices demonstrated a significant reduction in sIPSC frequency in HET organoids compared with WT (**Figure 5L–O**), indicating decreased inhibitory transmission.

To explore pharmacological intervention in a human cell-based model, we performed live calcium imaging in brain organoids combined with targeted GABA application (**Figure 5P**). Exogenous GABA (3 µM) inhibited neuronal activity in both WT and HET organoids (**Figure 5Q(i–iii), R**). Co-applying clonazepam (1 µM) with GABA (1 µM) significantly increased sIPSC frequency in both genotypes (**Figures S5I; 5S**). Calcium spike rates analysis under baseline, clonazepam, and washout conditions further showed that clonazepam normalized neuronal activity (**Figure 5T**). Although humans are *SCN2A* haploinsufficient, to further validate our findings, we generated homozygous (HOM) organoids, which exhibited complete loss of Na_V_1.2 expression (**Figure 5K**). Notably, enhancing GABA transmission also reduced neuronal activity in HOM organoids (**Figure 5R; S5J(i–iii)**).

Collectively, these findings indicate that *SCN2A* deficiency disrupts GABAergic gene expression and inhibitory transmission in both mouse and human cell-based models. Importantly, pharmacological enhancement of GABA_A_ receptor function not only reverses neuronal hyperexcitability and social deficits in Na_V_1.2-deficient mice but also restores inhibitory signaling and normalizes calcium dynamics in human brain organoids.

## DISCUSSION

In this study, we demonstrate that severe Na_V_1.2 deficiency leads to robust social deficits, altered synaptic transmission, and dysregulated *in vivo* neuronal activity, providing critical insights into *SCN2A*-associated disorders. Behaviorally, Na_V_1.2-deficient mice exhibited profound impairments across multiple social paradigms, including diminished exploration in the three-chamber tests and reduced reciprocal social interactions. These deficits are particularly striking in light of previous findings in heterozygous *Scn2a^+/–^* mice, which typically display normal, or even slightly enhanced, sociability^7–9^. This contrast underscores the pivotal role of Na_V_1.2 in regulating social function. Importantly, we demonstrate that the severe social deficits observed in our Na_V_1.2-deficient mice are reversible in adulthood through both global and circuit-specific restoration of *Scn2a* expression. At the cellular level, electrophysiological analyses indicate that Na_V_1.2 deficiency leads to an elevated E/I ratio. This imbalance is primarily attributable to a disproportionate reduction in inhibitory postsynaptic currents relative to excitatory inputs, which likely contributes to the observed *in vivo* basal neuronal hyperactivity and impaired SNR during social interactions. Consistent with this interpretation, pharmacological enhancement of GABA_A_ receptor function using clonazepam and AZD7325 alleviated social deficits, thereby identifying GABAergic dysregulation as a possible therapeutic target.

A key aspect of our findings lies in the distinction between canonical heterozygous knockout and gene-trap models of *Scn2a* deficiency. Early studies focusing on *Scn2a^+/–^* mice, necessitated by the perinatal lethality of complete knockouts (*Scn2a^−/−^*)^11^, which generally revealed minimal social abnormalities^7–9,42^. Similarly, mouse models carrying point mutations identified from human variants have reported only subtle or absent social impairments^29,43–45^. In contrast, our *Scn2a^gt/gt^*mice, with an approximate 70% reduction in Na_V_1.2 expression^20^, display severe social deficits (**Figures 1; S1–1**). This disparity suggests that mice are relatively resilient to moderate reductions in gene expression and that exceeding a critical limit (i.e., beyond a 50% reduction) may trigger a non-linear deterioration in neuronal function^10^. Indeed, while neurons with a 50% reduction in *Scn2a* maintain WT levels of excitability^28^, further reduction pushes neuronal properties past a tipping point, resulting in unexpected hyperexcitability^20,36,46–48^, a phenomenon that may underlie the seizure phenotypes observed in children with *SCN2A* protein-truncating variants and whole gene deletions^49^. Another notable difference between *Scn2a^+/−^* mice and *Scn2a^gt/gt^*mice is the E/I imbalance. A previous study in adult *Scn2a^+/–^*mice reported a decrease in mEPSC frequency without significant changes in mIPSC frequency, suggesting a net reduction in mE/I ratio^28^. In contrast, our *Scn2a^gt/gt^* mice displayed a reduction in mEPSC frequency^48^ coupled with an even greater decrease in mIPSC frequency, resulting in an elevated mE/I ratio (**Figure 1**). This pattern, consistent with findings observed across multiple ASD models^15^, likely underpins the profound social deficits correlated to the increased E/I imbalance and may represent a converging mechanism underlying autistic-like phenotypes.

Our gene-trap strategy was designed to model the complete loss-of-function (LoF) of *SCN2A*, including the nonsense mutation *SCN2A-C959X*, which introduces a premature stop codon and triggers nonsense-mediated decay^6,41^. These nonconducting variants lack dominant-negative effects^50^ and result in haploinsufficiency, where the 50% protein level from the unaffected allele is insufficient to prevent phenotypic manifestations^48^. While *SCN2A* missense mutations, particularly those with dominant-negative effects^45,51^, often produce mutant proteins that aggregate and sequester WT Na_V_1.2, further reducing functional protein levels below 50%. Our homozygous *SCN2A-C959X* mutation, representing an extreme condition of dominant-negative disruption, confirms the neuronal phenotypes observed in human brain organoids carrying heterozygous *SCN2A-C959X* mutation and aligns with our gene-trap mice (**Figure 5**). To our knowledge, this is the first *Scn2a*-deficient mouse model to robustly exhibit social deficits and impaired GABAergic transmission, effectively recapitulating the human cell-based models and patient behavioral phenotypes. Our findings establish a valuable platform for investigating both nonsense and missense LoF *SCN2A* mutations and for testing therapeutic interventions for social impairments.

Our study also reveals that the social behavior in Na_V_1.2-deficient mice is remarkably dynamic. Notably, even in adulthood, social deficits can be rescued by restoring *Scn2a* expression, and conversely, reintroducing *Scn2a* null reverses these gains (**Figure 3**). This dynamic modulation of sociability by altering *Scn2a* expression is unexpected given the prevailing view that social behavior is governed by complex, multi-regional networks involving numerous neurotransmitters and receptors^52^. Instead, our findings indicate a surprisingly direct relationship between Na_V_1.2 expression and social behavior, with significant implications for understanding the molecular underpinnings of sociability.

At the circuit level, our results underscore the critical role of striatal projection networks in mediating social behavior. Although social behaviors are traditionally considered to be orchestrated by distributed neural circuits, the striatum, which integrates diverse inputs, appears to serve as a key node for social processing^53,54^. Notably, our data indicate that restoring Na_V_1.2 in NAc-projecting neurons, rather than in striatal neurons *per se*, is sufficient to rescue social deficits (**Figure 4**), suggesting that upstream projection neurons are primary drivers of sociability. This observation implies that the net synaptic afferents, as reflected in the E/I balance, may underlie the social impairments associated with *Scn2a* deficiency. Among these upstream circuits, the mPFC-NAc projection emerges as particularly critical^55^. Future studies will explore whether targeted inhibition of excitatory mPFC neurons projecting to the striatum can alleviate social deficits, thereby reinforcing the importance of this circuit. Although other circuits, such as the mPFC-amygdala^56^ implicated in *ArpC3*^57^ and *Pten^+/-^*^58^ mice and the mPFC-posterior paraventricular nucleus of the thalamus (pPVT)^59^, have also been associated with various aspects of social behavior, further research is needed to dissect their specific contributions in the context of Na_V_1.2 deficiency.

The NAc functions as a center node in brain circuits governing social behaviors^13,14^. In our study, *Scn2a* deficiency leads to basal neuronal hyperactivity (**Figure 2**), similar to the hyperactivity seen in a *Shank3* mouse model with social impairments^60^. We considered this behaviorally irrelevant excessive basal activity as “noise” and demonstrated that normalizing it, by *Scn2a* restoration and enhanced GABAergic transmission, rescued social interactions in *Scn2a*-deficient mice. Notably, Zhao et al. reported that nonspecific inhibition of NAc core neurons similarly improved social behavior in a *Cntnap*^2*-/-*^ mouse model^61^. Moreover, using an algorithm for social neural ensembles in the mPFC^62^, we identified behaviorally tuned social-ON and social-OFF neurons. The predominance of social-ON neurons aligns with previous sociability assays linking social engagement to MSNs^61^. We observed that *Scn2a* restoration selectively enhanced social-ON signaling by reducing basal noise, whereas further *Scn2a* suppression exacerbated aberrant neuronal activity and disrupted social-OFF signal encoding (**Figure S4–2M**). Given that D1-MSNs are less excitable than D2-MSNs^63,64^ and typically drive opposing behaviors^65^, our data suggest that social-ON neurons might correspond to D1-MSNs, and while social-OFF neurons might correspond to D2-MSNs, as supported by reciprocal effects following their inhibition^61^.

In summary, our findings reveal that severe Na_V_1.2 deficiency produces profound and reversible social deficits, underpinned by disproportionate reductions in excitatory and inhibitory synaptic transmission. We demonstrate a direct, dose-dependent relationship between *Scn2a* expression and sociability, whereby a ∼70% reduction in Na_V_1.2 leads to an elevated E/I ratio, increased noisy basal activity, and impaired neuronal coding, while restoration of *Scn2a* or pharmacological enhancement of GABA_A_ receptor function reverses these deficits. Collectively, our work provides a comprehensive exploration, from molecular and cellular mechanisms to neural circuits, of the pathophysiology underlying social impairments in *SCN2A*-associated disorders. These insights lay a robust foundation for the development of targeted therapeutic interventions aimed at normalizing synaptic function to ameliorate social impairments.

## ACKNOWLEDGMENTS

We sincerely thank Dr. Julie M. J. Fabre from the UCL Queen Square Institute of Neurology for her invaluable assistance in Neuropixels data analysis. Additionally, we thank Dr. Shulan Xiao from the Department of Biomedical Engineering at Purdue University for calcium imaging analysis. This work is supported by Purdue startup funding, Ralph W. and Grace M. Showalter Research Trust, and Purdue Big Idea Challenge 2.0 on Autism to Y.Y. Research reported in this publication was also supported by the National Institute of Neurological Disorders and Stroke of the National Institutes of Health under Award Number R01NS117585 and R01NS123154 to Y.Y. The authors gratefully acknowledge support from the Purdue University Institute for Drug Discovery and Institute for Integrative Neuroscience for additional funding support. M.E. is supported by the National Science Foundation (NSF) Graduate Research Fellowship Program (GRFP) (DGE-1842166). X.C. is supported by the AES Postdoctoral Research Fellowship. Yang lab is grateful to the *FamilieSCN2A* foundation for the Action Potential Grant support to M.E. X.C. and J.Z. This project was funded, in part, with support from the Indiana Spinal Cord & Brain Injury Research Fund from the Indiana State Department of Health, and the Indiana Clinical and Translational Sciences Institute (CTSI) funded, in part by Award Number UL1TR002529 from the National Institutes of Health, National Center for Advancing Translational Sciences, Clinical and Translational Sciences Award. The Yang lab appreciates the bioinformatics support of the Collaborative Core for Cancer Bioinformatics (C^3^B) with support from the Indiana University Simon Comprehensive Cancer Center (Grant P30CA082709), Purdue University Center for Cancer Research (Grant P30CA023168), and Walther Cancer Foundation. The content is solely the responsibility of the authors and does not necessarily represent the official views of the Indiana State Department of Health, National Institutes of Health, or the National Science Foundation.

## AUTHOR CONTRIBUTIONS

J.Z., M.E., X.C., Y.Y. designed the experiments. J.Z., M.E., X.C., Y.Z., S.K., B.A.D., K.H., H.P.N., Y.S., S.L., J.W., Z.Q., K.W.W., Z.Z., T.X., M.S.H., M.I.O.A., Y.E.Y., performed the experiments and analyzed the data. N.A.L., W.A.K., W.C.S. participated in data analysis and experimental design. Y.Y. supervised the project. J.Z., M.E., X.C., and Y.Y. wrote the paper with input from all authors.

## Supplemental Information

**Figure S1–1.**
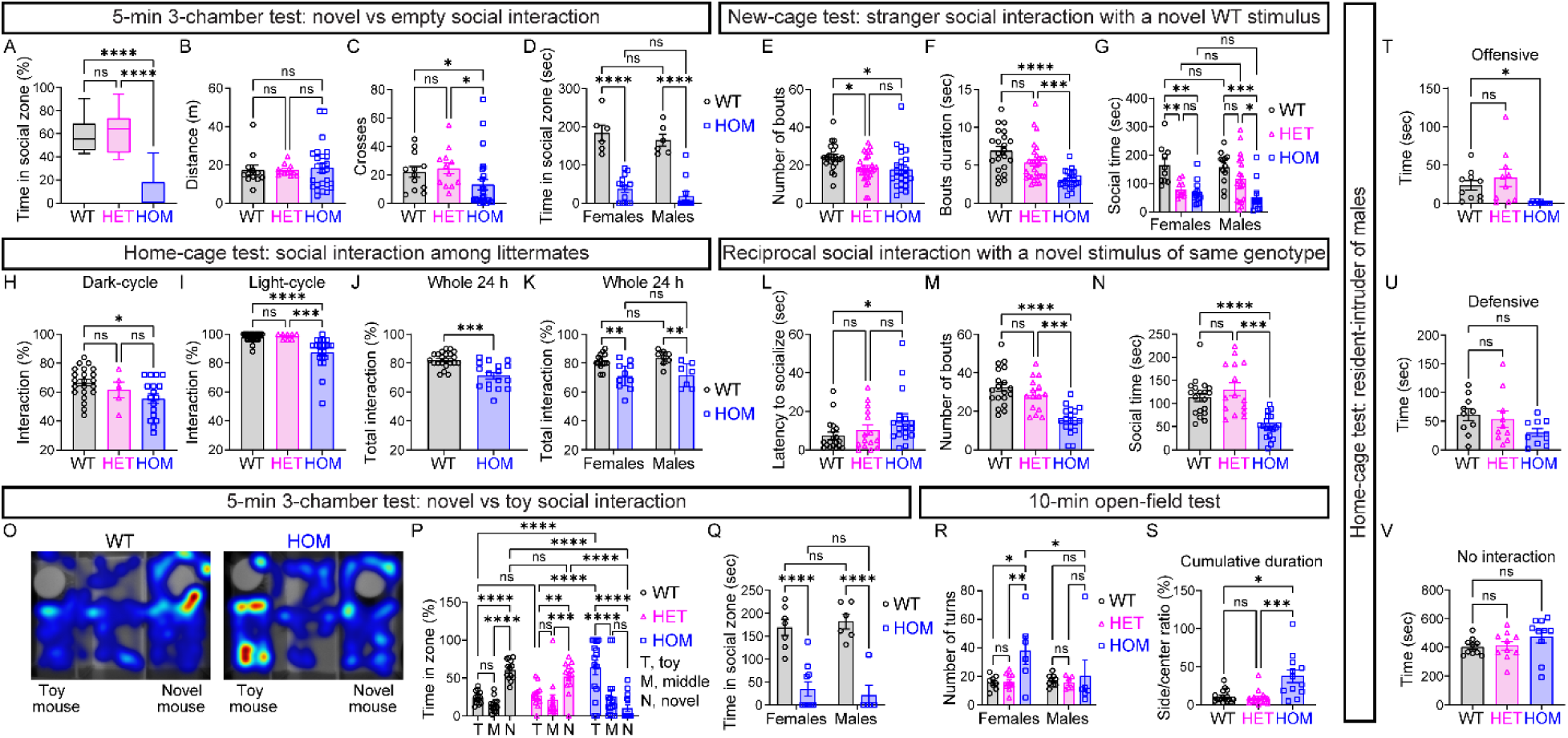
*Scn2a* deficiency leads to social deficits across multiple behavior assays. Related to Figure 1. (A–D) Three-chamber sociability test (novel mouse versus empty) of *Scn2a^+/+^*(WT, black), *Scn2a^+/gt^* (HET, magenta), and *Scn2a^gt/gt^* (HOM, blue) mice, comparing time spent in the social zone, distance, and crosses traveled. n = 12 WT, n = 12 HET, and n = 26 HOM mice. One-way ANOVA with Tukey’s multiple comparisons (A), Kruskal-Wallis with Dunn’s multiple comparisons (B, C), and two-way ANOVA with Fisher’s LSD test (D): ns, not significant; *p < 0.05; ****p < 0.0001. WT = Wild Type, HET = Heterozygous, HOM = Homozygous. (E–G) New-cage test examining social interaction bouts, duration, and social time with a novel WT stimulus. n = 22 WT, n = 30 HET, and n = 25 HOM mice. One-way ANOVA with Tukey’s multiple comparisons (E), Kruskal-Wallis with Dunn’s multiple comparisons (F), and two-way ANOVA with Sidak’s multiple comparisons (G): ns, not significant; *p < 0.05; **p < 0.01; ***p < 0.001; ***p < 0.0001. (H–K) Home-cage social interactions between littermates during the dark cycle, light cycle, and 24-hour periods. One-way ANOVA with Tukey’s multiple comparisons (H), Kruskal-Wallis with Dunn’s multiple comparisons (I), Welch’s t test (J), and two-way ANOVA with Fisher’s LSD test (K): ns, not significant; *p < 0.05; **p < 0.01; ***p < 0.001; ****p < 0.0001. (L–N) Reciprocal social interactions with a novel stimulus of the same genotype. n = 19 WT pairs, n = 15 HET pairs, and n = 18 HOM pairs of mice. Kruskal-Wallis with Dunn’s multiple comparisons (L), one-way ANOVA with Tukey’s multiple comparisons (M), and Brown-Forsythe and Welch one-way ANOVA with Dunnett T3’s multiple comparisons (N): ns, not significant; *p < 0.05; ***p < 0.001; ****p < 0.0001. (O–Q) Three-chamber sociability test assessing preference for a novel live mouse versus a toy mouse. (O) Representative heatmaps of tracking trajectories for a single social preference (novel mouse versus toy) trial of WT and HOM mice (top view of set up). Warm colors (red, orange, yellow) represent more frequented areas whereas cool colors (green and blue) are less frequented areas. (P) Percentage of cumulative time spent in each zone (toy, middle chamber, or novel mouse). (Q) Cumulative time spent in the chamber with the novel mouse. n = 13 WT (7 females + 6 males), n = 13 HET (7 females + 6 males), and n = 15 HOM mice (10 females + 5 males). Two-way ANOVA with Fisher’s LSD test: ns, not significant; **p < 0.01; ***p < 0.001; ****p < 0.0001. (R–S) Open-field test for turns and side preference. n = 17 WT (8 females + 9 males), n = 17 HET (11 females + 6 males), and n = 12 HOM mice (6 females + 6 males). Two-way ANOVA with Tukey’s multiple comparisons (R) and Kruskal-Wallis with Dunn’s multiple comparisons (S): ns, not significant; *p < 0.05; **p < 0.01; ***p < 0.001. (T–V) Home-cage resident-intruder test measuring offensive, defensive, and no interaction times. n = 10 WT, n = 10 HET, and n = 10 HOM mice. Brown-Forsythe and Welch one-way ANOVA with Dunnett T3’s multiple comparisons (T) and one-way ANOVA with Dunnett’s multiple comparisons (U, V): ns, not significant; *p < 0.05. Data are represented as mean ± SEM.

**Figure S1–2.**
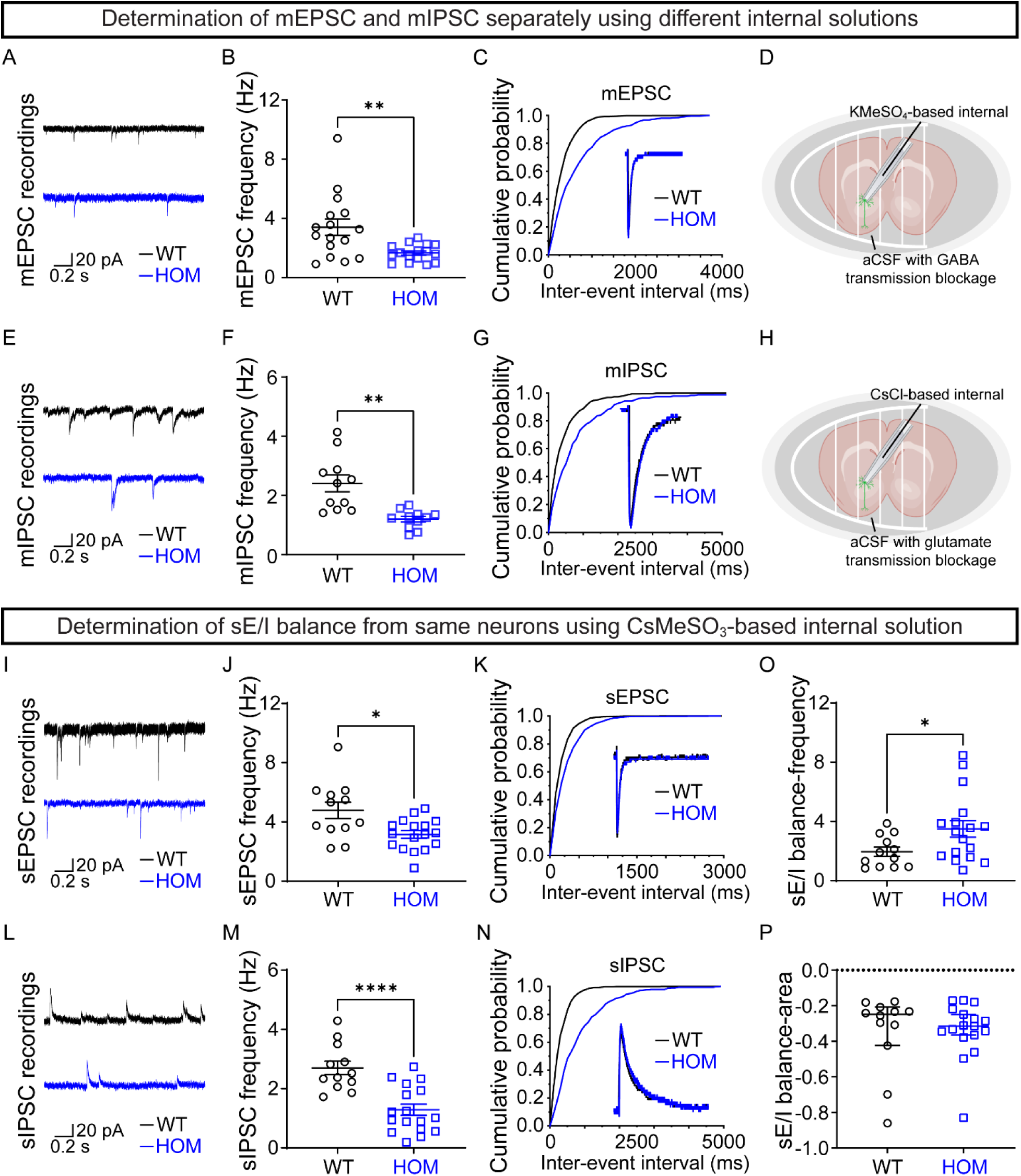
*Scn2a* deficiency renders a greater reduction in GABA transmission onto MSN in the NAc core. Related to Figure 1. (A) Example traces of mEPSCs recorded on MSNs from *Scn2a^+/+^* (WT, upper, black) and *Scn2a^gt/gt^* (HOM, lower, blue) mice. (B) mEPSC frequency (n = 16 cells from 3 WT mice and n = 16 cells from 3 HOM mice). Unpaired Welch’s t test: **p < 0.01. (C) Cumulative probability of mEPSC inter-event intervals. Inset: representative averaged mEPSCs. (D) Schematic of experiments in (A) to (C) using KMeSO_4_-based internal solution. (E) Example traces of mIPSCs recorded on MSNs from WT (upper, black) and HOM (lower, blue) mice. (F) mIPSC frequency (n = 11 cells from 3 WT mice and n = 11 cells from 3 HOM mice). Unpaired Welch’s t test: **p < 0.01. (G) Cumulative probability of mIPSCs inter-event intervals. Inset: representative averaged mIPSCs. (H) Schematic of experiments in (E) to (G) using CsCl-based internal solution. (I) Example traces of sEPSCs recorded on MSNs from WT (upper, black) and HOM (lower, blue) mice using CsMeSO_3_-based internal solution. (J) sEPSC frequency (n = 12 cells from 6 WT mice and n = 17 cells from 7 HOM mice). Unpaired Welch’s t test: *p < 0.05. (K) Cumulative probability of sEPSC inter-event intervals. Inset: representative averaged mEPSCs. (L) Example traces of mIPSCs recorded on MSNs from WT (upper, black) and HOM (lower, blue) mice using CsMeSO_3_-based internal solution. (M) sIPSC frequency (n = 12 cells from 6 WT mice and n = 17 cells from 7 HOM mice). Unpaired t test: ****p < 0.0001. (N) Cumulative probability of mIPSCs inter-event intervals. Inset: representative averaged sIPSCs. (O) sEPSC/sIPSC ratio calculated with frequency. Student’s t test: *p < 0.05. (P) sEPSC/sIPSC ratio calculated with area under the curve. Student’s t test: ns, not significant. Data were shown as mean ± SEM.

**Figure S2.**
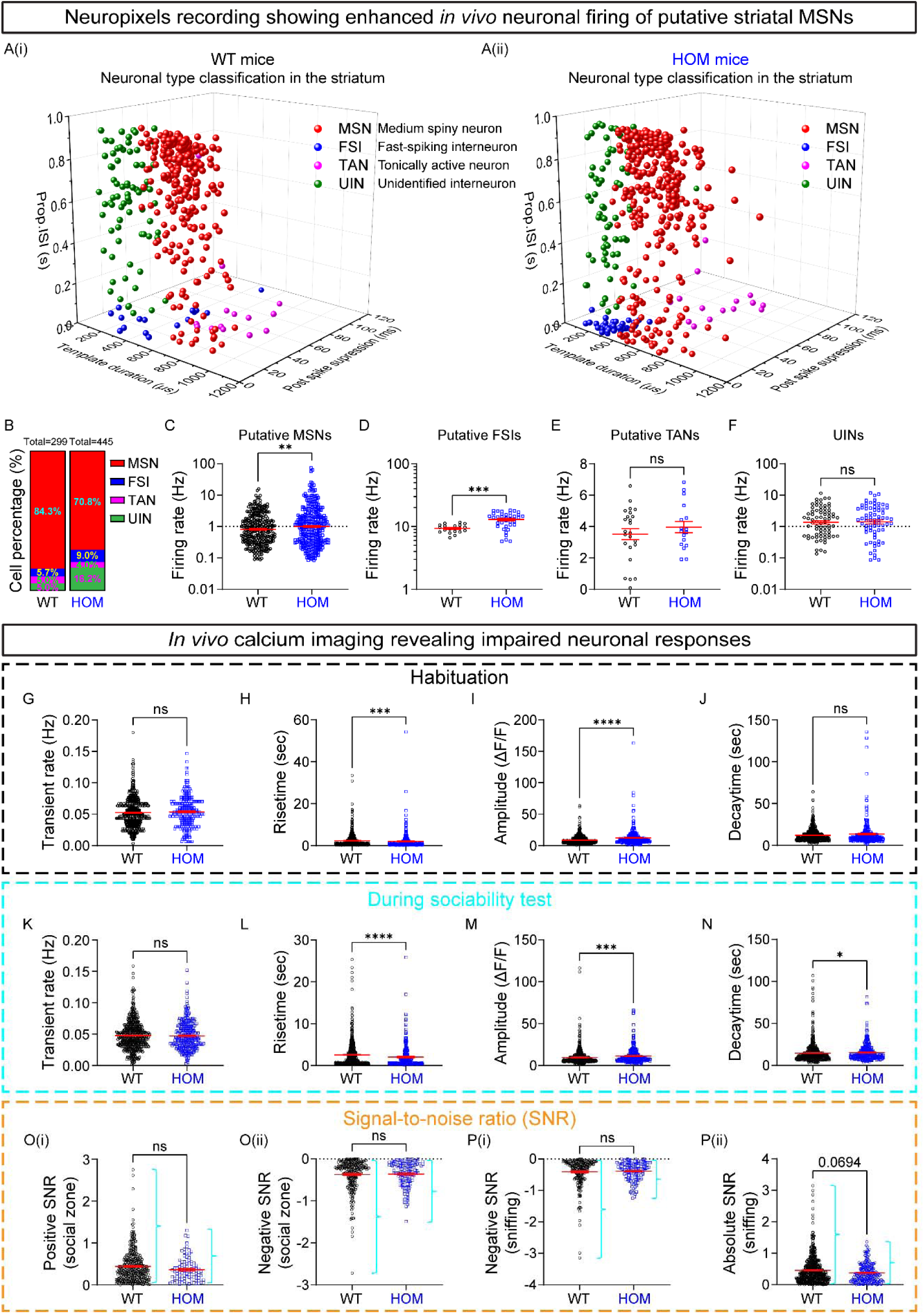
Elevated basal *in vivo* neuronal activity and impaired SNR in *Scn2a* deficient mice. Related to Figure 2. (A–F) Neuropixels recordings of neuronal firing in the striatum. (A) Neuronal subtype classification in the striatum for *Scn2a^+/+^* (WT, A(i)) and *Scn2a^gt/gt^*(HOM, (A(ii)): putative medium spiny neurons (MSNs), fast-spiking interneurons (FSIs), tonically active neurons (TANs), and unidentified interneurons (UINs). (B) Percentage distribution of cell types in WT and HOM mice. (C–F) Firing rates of neuronal subtypes in the striatum. HOM mice show significantly increased firing rates in putative MSNs (C) and FSIs (D), with no significant differences in TANs (E) or UINs (F). n = 288 MSNs, 18 FSIs, 24 TANs, and 73 UINs from 8 WT mice; and n = 314 MSNs, 39 FSIs, 17 TANs, and 71 UINs from 8 HOM mice. Linear mixed-effects model fit by maximum likelihood: ns, not significant; **p < 0.01; ***p < 0.001. Data are presented as mean ± SEM. (G–J) Calcium transient properties during habituation. HOM mice exhibit a comparable Ca^2+^ transient rate and decaytime but display a faster risetime and increased amplitude compared with WT mice, indicating elevated basal neuronal activity. n = 623 cells from 3 WT mice and n = 307 cells from 5 HOM mice. Unpaired Mann-Whitney *U* test: ns, not significant; ***p < 0.001; ****p < 0.0001. (K–N) Calcium transient properties during sociability tests. n = 623 cells from 3 WT mice and n = 307 cells from 5 HOM mice. Unpaired Mann-Whitney *U* test: ns, not significant; *p < 0.05; ***p < 0.001; ****p < 0.0001. (O–P) Signal-to-noise ratio (SNR) analysis during social interaction tests. n = 626 (380 positive and 246 negative, social zone; 236 negative, sniffing) cells from 3 WT mice and n = 228 (93 positive and 135 negative, social zone; 125 negative, sniffing) cells from 3 HOM mice. Unpaired Mann-Whitney *U* test: ns, not significant. Data are presented as mean ± SEM.

**Figure S3–1.**
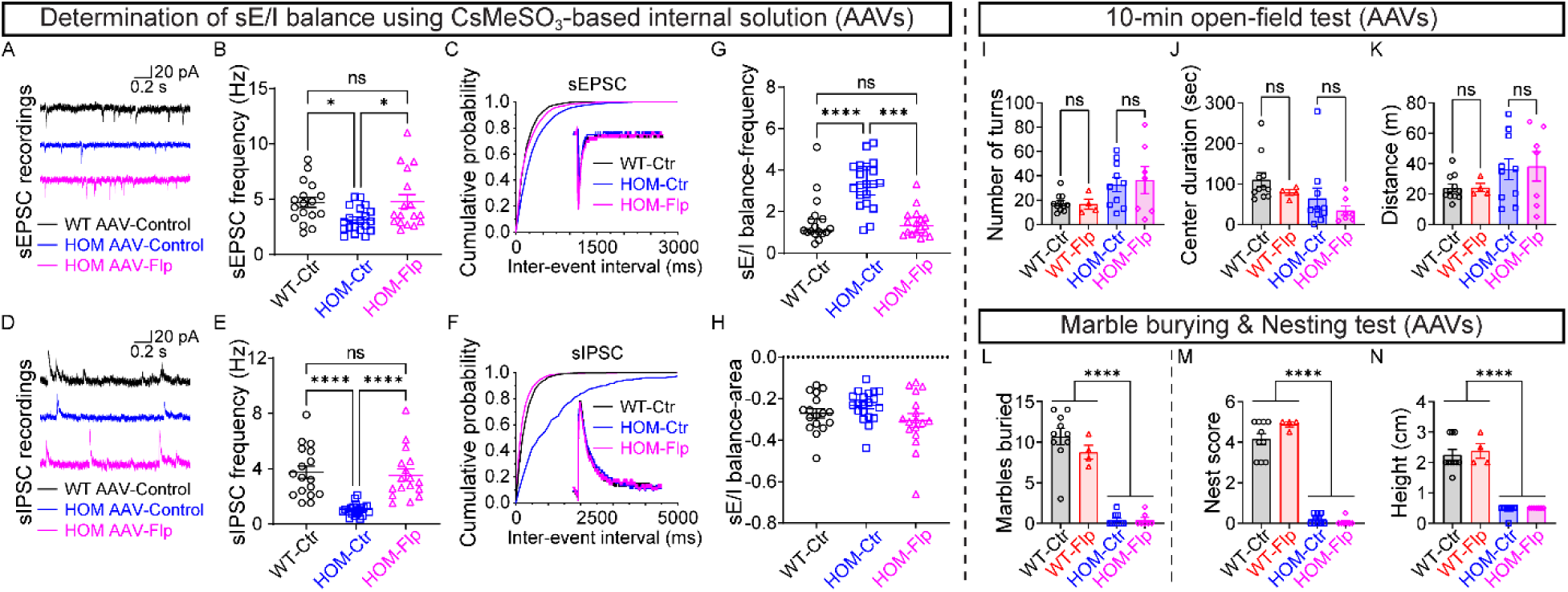
Global restoration of *Scn2a* with AAV-PHP.eB-Flp reverses sE/I ratio but not anxiety or other behaviors associated with autism in adult Na_V_1.2-deficient mice. Related to Figure 3. (A–C) Analysis of sEPSCs recorded on MSNs from *Scn2a^+/+^* (WT) mice transduced with AAV-Control (n = 17 cells from 6 mice, black), *Scn2a^gt/gt^* (HOM) mice transduced with AAV-Control (n = 20 cells from 7 mice, blue), and HOM mice transduced with AAV-Flp (n = 17 cells from 6 mice, magenta). (A) Representative sEPSC traces, (B) frequency, and (C) cumulative probability of inter-event intervals (Inset: representative averaged sEPSCs). One-way ANOVA with Tukey’s multiple comparisons: ns, not significant; *p < 0.05. (D–F) Analysis of sIPSCs recorded on MSNs from WT mice transduced with AAV-Control (n = 17 cells from 6 mice, black), HOM mice transduced with AAV-Control (n = 20 cells from 7 mice, blue), and HOM mice transduced with AAV-Flp (n = 17 cells from 6 mice, magenta). (D) Representative sIPSC traces, (E) frequency, and (F) cumulative probability of inter-event intervals (Inset: representative averaged sIPSCs). Brown-Forsythe and Welch one-way ANOVA with Dunnett T3’s multiple comparisons: *p < 0.05. (G, H) The sE/I ratio by frequency (M) and area (N) highlighting restored sE/I balance in HOM mice transduced with AAV-Flp. Kruskal-Wallis with Dunn’s multiple comparisons (G) and one-way ANOVA with Tukey’s multiple comparisons (H): ns, not significant; *p < 0.05. (I–K) Open-field test results showing (I) number of turns, (J) center duration, and (K) total distance traveled. n = 11 WT mice transduced with AAV-Control, n = 4 WT mice transduced with AAV-Flp, n = 10 HOM mice transduced with AAV-Control, and n = 7 HOM mice transduced with AAV-Flp. One-way ANOVA with Bonferroni’s multiple comparisons: ns, not significant. (L) Marble-burying test showing the number of marbles buried by each group. n = 10 WT mice transduced with AAV-Control, n = 4 WT mice transduced with AAV-Flp, n = 10 HOM mice transduced with AAV-Control, and n = 7 HOM mice transduced with AAV-Flp. One-way ANOVA with Bonferroni’s multiple comparisons: ****p < 0.0001. (M, N) Nest-building behaviors showing (M) nest score and (N) nest height. n = 10 WT mice transduced with AAV-Control, n = 4 WT mice transduced with AAV-Flp, n = 10 HOM mice transduced with AAV-Control, and n = 7 HOM mice transduced with AAV-Flp. One-way ANOVA with Bonferroni’s multiple comparisons: ****p < 0.0001. Data were shown as mean ± SEM.

**Figure S3–2.**
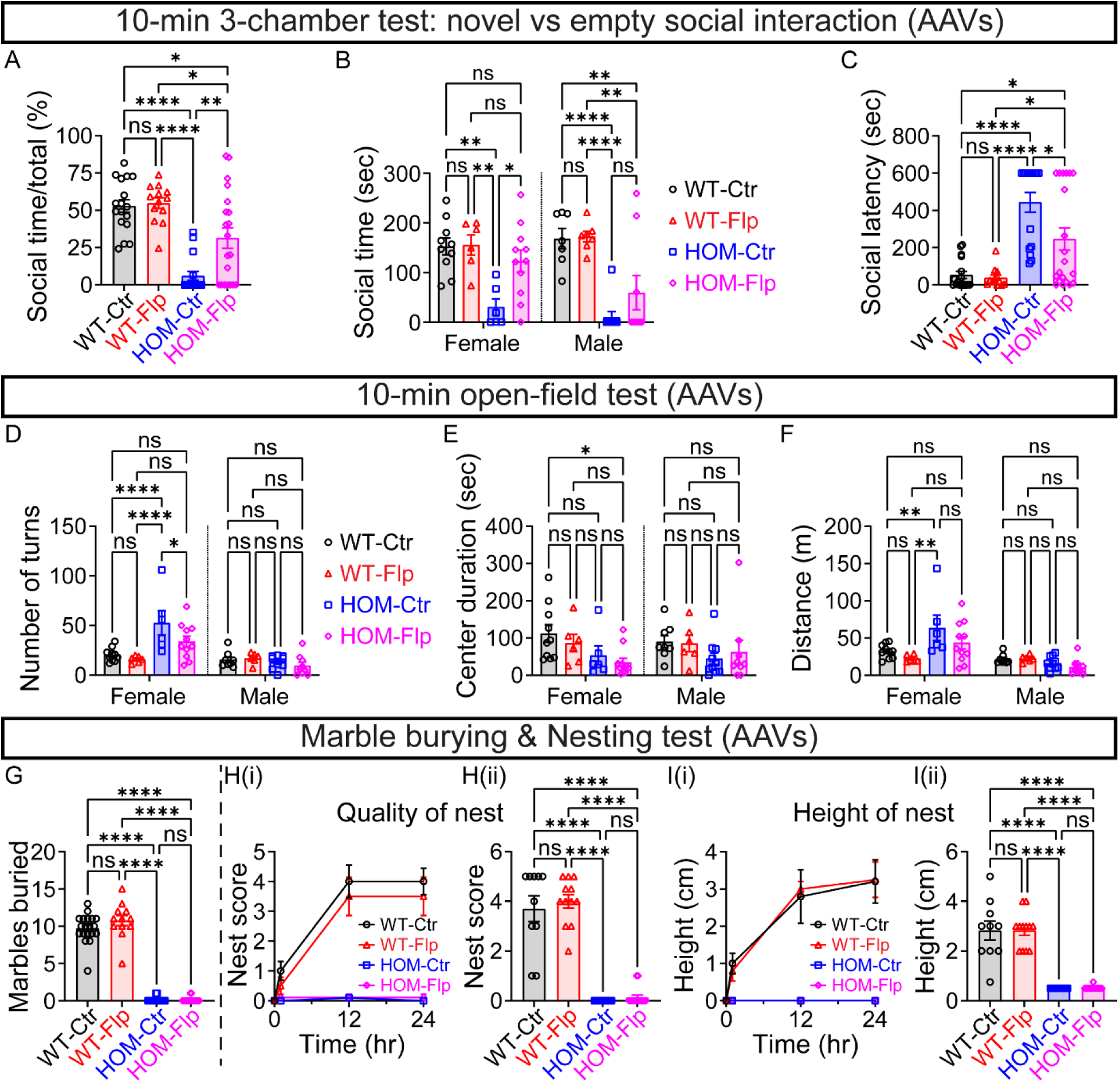
Global restoration of *Scn2a* with a lower AAV-PHP.eB-Flp dose (2×10^11^ genome copies (GC) per mouse) alleviates social impairments in female mice, but not anxiety or other behaviors associated with autism. Related to Figure 3. (A–C) Quantification of social interaction behaviors in the three-chamber sociability test assessing social preference (novel mouse versus empty). (A) Social interaction percentage (social interaction/ total time), (B) total social interaction time stratified by sex, and (C) latency to novel stimulus. n = 17 (10 females + 7 males) WT mice transduced with AAV-Control, n = 13 (6 females + 7 males) WT mice transduced with AAV-Flp, n = 16 (6 females + 10 males) *Scn2a^gt/gt^* (HOM) mice transduced with AAV-Control, and n = 20 (11 females + 9 males) HOM mice transduced with AAV-Flp. One-way ANOVA with Bonferroni’s multiple comparisons (A), two-way ANOVA with Tukey’s multiple comparisons (B), and one-way ANOVA with Sidak’s multiple comparisons (C): ns, not significant; *p < 0.05; **p < 0.01; ****p < 0.0001. (D–F) Open-field test results showing number of turns (D), time spent in the center of the open field (E), and total distance traveled (F), stratified by sex. n = 18 (10 females + 8 males) WT mice transduced with AAV-Control, n = 12 (6 females + 6 males) WT mice transduced with AAV-Flp, n = 16 (6 females + 10 males) HOM mice transduced with AAV-Control, and n = 20 (11 females + 9 males) HOM mice transduced with AAV-Flp. Two-way ANOVA with Tukey’s (D, F) and Bonferroni’s (E) multiple comparisons: ns, not significant; *p < 0.05; **p < 0.01; ****p < 0.0001. (G) Marble-burying test showing the number of marbles buried by each group. n = 19 WT mice transduced with AAV-Control, n = 12 WT mice transduced with AAV-Flp, n = 16 HOM mice transduced with AAV-Control, and n = 20 HOM mice transduced with AAV-Flp. One-way ANOVA with Sidak’s multiple comparisons: ns, not significant; ****p < 0.0001. (H, I) Nest-building behaviors over a 24-hour period. (H) Quality of nest score: (H(i)) timeline of nest scores at 12 and 24 hours and (H(ii)) quantification of nest quality at 24 hours. (I) Nest height measurements: (I(i)) timeline of nest height and (I(ii)) quantification at 24 hours. n = 10 WT mice transduced with AAV-Control, n = 12 WT mice transduced with AAV-Flp, n = 14 HOM mice transduced with AAV-Control, and n = 15 HOM mice transduced with AAV-Flp. One-way ANOVA with Bonferroni’s multiple comparisons: ns, not significant; ****p < 0.0001. Data are represented as mean ± SEM.

**Figure S4–1.**
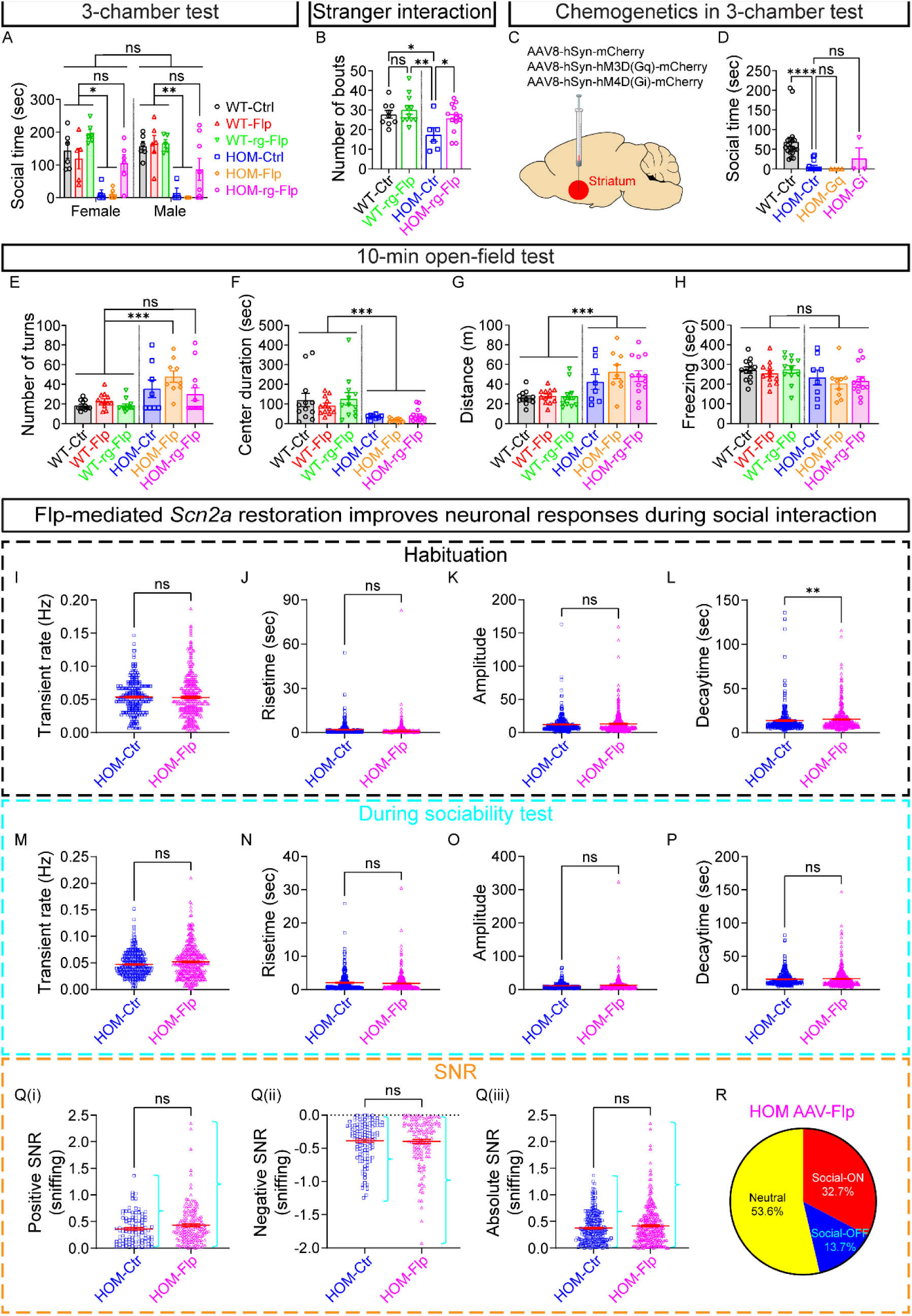
Restoration of *Scn2a* expression in NAc-projecting neurons rescues social deficits but not anxiety-like behaviors in adult Na_V_1.2-deficient mice. Related to Figure 4. (A) Three-chamber test showing social time spent in the chamber with the novel mouse, stratified by sex (novel mouse versus empty). n = 12 (6 females + 6 males) WT mice transduced with AAV-Control (black), n = 11 (5 females + 6 males) WT mice transduced with AAV-Flp (red), n = 11 (6 females + 5 males) WT mice transduced with retrograde AAV-Flp (green), n = 9 (5 females + 4 males) *Scn2a^gt/gt^* (HOM) mice transduced with AAV-Control (blue), n = 9 (6 females + 3 males) HOM mice transduced with AAV-Flp (orange), and n = 14 (6 females + 8 males) HOM mice transduced with retrograde AAV-Flp (magenta). Two-way ANOVA with Tukey’s multiple comparisons: ns, not significant; *p < 0.05; **p < 0.01. (B) Number of interaction bouts during the stranger interaction test. n = 9 WT mice transduced with AAV-Control (black), n = 11 WT mice transduced with retrograde AAV-Flp (green), n = 6 HOM mice transduced with AAV-Control (blue), and n = 14 HOM mice transduced with retrograde AAV-Flp (magenta). Kruskal-Wallis with uncorrected Dunn’s multiple comparisons: ns, not significant; *p < 0.05; **p < 0.01. (C) Schematic illustrating targeted viral delivery to striatal neurons for chemogenetic manipulation using AAV8-hSyn-hM3D(Gq)-mCherry or AAV8-hSyn-hM4D(Gi)-mCherry, which provide excitatory or inhibitory modulation, respectively. A control group was injected with AAV8-hSyn-mCherry lacking the DREADD cassette. (D) Three-chamber sociability test (novel mouse versus empty) showing social time of the mice transduced with Gq/Gi-DREADD targeting striatal neurons only. n = 19 WT mice transduced with AAV-Control (black), n = 16 HOM mice transduced with AAV-Control (blue), n = 4 HOM mice transduced with AAV-Gq (orange), and n = 3 HOM mice transduced with AAV-Gi (magenta). One-way ANOVA Dunnett’s multiple comparisons: ns, not significant; ****p < 0.0001. (E–H) Open-field test results assessing general locomotor and anxiety-like behaviors. (E) Number of turns, (F) center duration, (G) distance traveled, and (H) freezing time. n = 12 WT mice transduced with AAV-Control (black), n = 12 WT mice transduced with AAV-Flp (red), n = 12 WT mice transduced with retrograde AAV-Flp (green), n = 8 HOM mice transduced with AAV-Control (blue), n = 9 HOM mice transduced with AAV-Flp (orange), and n = 13 HOM mice transduced with retrograde AAV-Flp (magenta). One-way ANOVA with Bonferroni’s multiple comparisons: ns, not significant; ***p < 0.001. (I–P) Analysis of neuronal responses during habituation (I–L) and sociability tests (M–P) in HOM mice transduced with AAV-Control (blue), and HOM mice transduced with AAV-Flp (magenta). (I, M) Transient rate (Hz), (J, N) rise time, (K, O) amplitude, and (L, P) decay time of neuronal calcium spikes during the respective behavioral paradigms. n = 307 cells from 5 HOM mice transduced with AAV-Control and n = 360 cells from 8 HOM mice transduced with retrograde AAV-Flp. Unpaired Mann-Whitney *U* test: ns, not significant; **p < 0.01. (Q) Signal-to-noise ratio (SNR) analysis. Positive SNR (Q(i), negative SNR (Q(ii)), and absolute SNR (Q(iii)) during sniffing events. n = 228 (103 positive sniffing + 125 negative sniffing) cells from 3 WT mice; and n = 296 (167 positive sniffing + 129 negative sniffing) cells from 6 HOM mice. Unpaired Mann-Whitney *U* test: ns, not significant. (R) Pie charts showing the distribution of social-ON, social-OFF, and neutral neurons from HOM mice transduced with AAV-Flp: social-ON (32.7%), social-OFF (13.7%), and neutral (53.6%). Data are represented as mean ± SEM.

**Figure S4–2.**
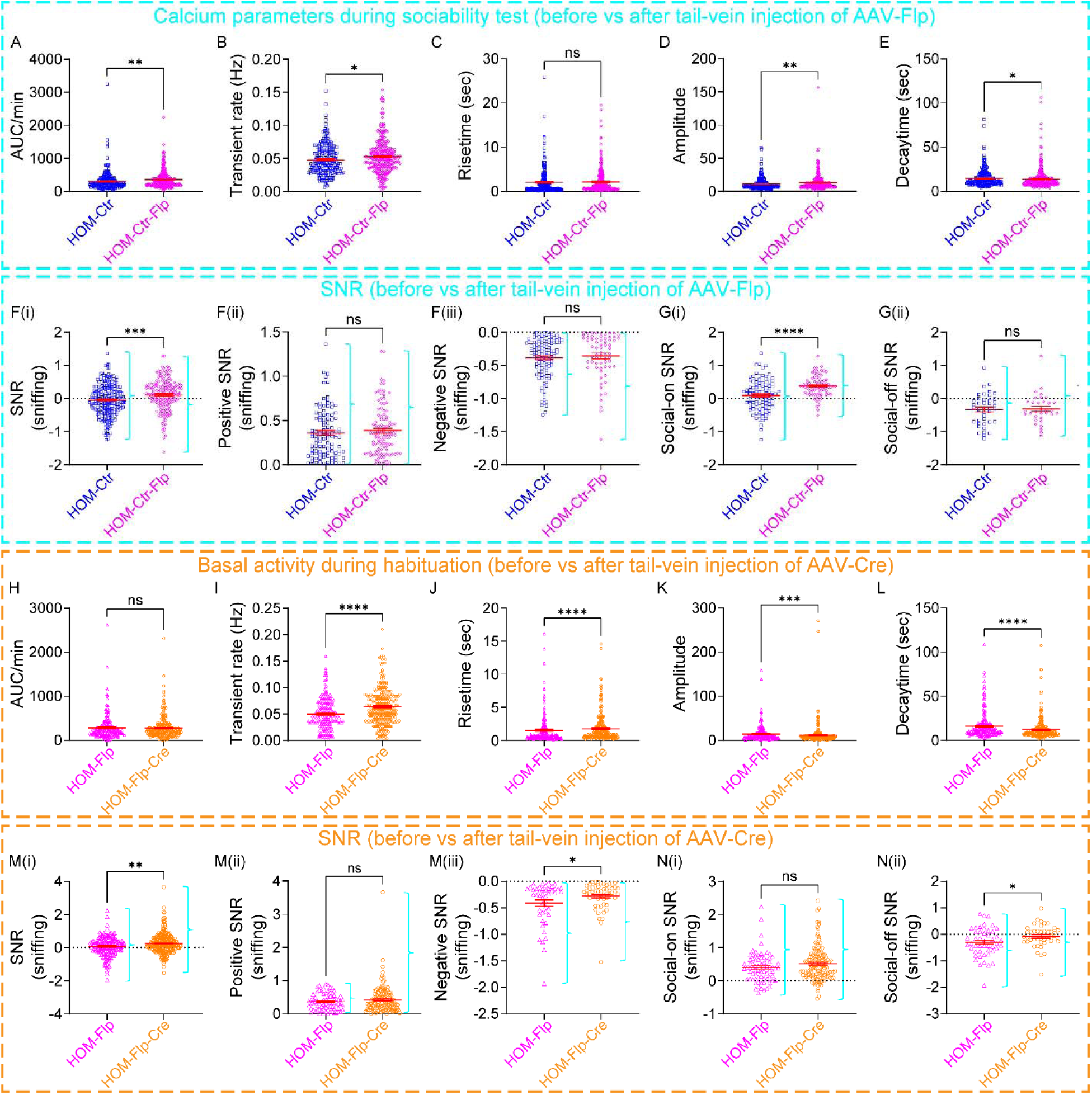
Calcium dynamics and SNR disturbance in response to bidirectional regulation of *Scn2a* in adult Na_V_1.2-deficient mice during behavioral paradigms. Related to Figure 4. (A–E) Calcium activity parameters during the sociability test, comparing HOM-Ctr mice before and after tail-vein injection of AAV-Flp. AUC (A), transient rate (B), rise time (C), amplitude (D), and decay time (E) of neuronal calcium spikes. n = 4 HOM mice (272 cells before + 328 cells after). Unpaired Mann-Whitney *U* test: ns, not significant; *p < 0.05; **p < 0.01. (F–G) SNR analysis in the sociability test, comparing HOM-Ctr mice before and after tail-vein injection of AAV-Flp. Overall SNR (F(i)), positive SNR (F(ii)), negative SNR (F(iii)), social-ON SNR(G(i)), and social-OFF SNR (G(i)) during sniffing events. Unpaired Mann-Whitney *U* test (F(i)–F(iii)), Welch’s test (G(i)), and t test (G(ii)): ns, not significant; ***p < 0.001; ****p < 0.0001. (H–L) Basal calcium activity parameters during habituation, comparing HOM-Flp mice before and after tail-vein injection of AAV-Cre. AUC (H), transient rate (I), rise time (J), amplitude (K), and decay time (L) of neuronal calcium spikes. n = 5 HOM mice (224 cells before + 289 cells after). Unpaired Mann-Whitney *U* test: ns, not significant; ***p < 0.001; ****p < 0.0001. (M–N) SNR analysis in the sociability test, comparing HOM-Flp mice before and after tail-vein injection of AAV-Cre. Overall SNR (M(i)), positive SNR (M(ii)), negative SNR (M(iii)), social-ON SNR(N(i)), and social-OFF SNR (N(i)) during sniffing events. Unpaired Mann-Whitney *U* test (M(i), M(ii), and N(i)), t test (M(iii), N(ii)): ns, not significant; *p < 0.05; **p < 0.01. Data are represented as mean ± SEM.

**Figure S5.**
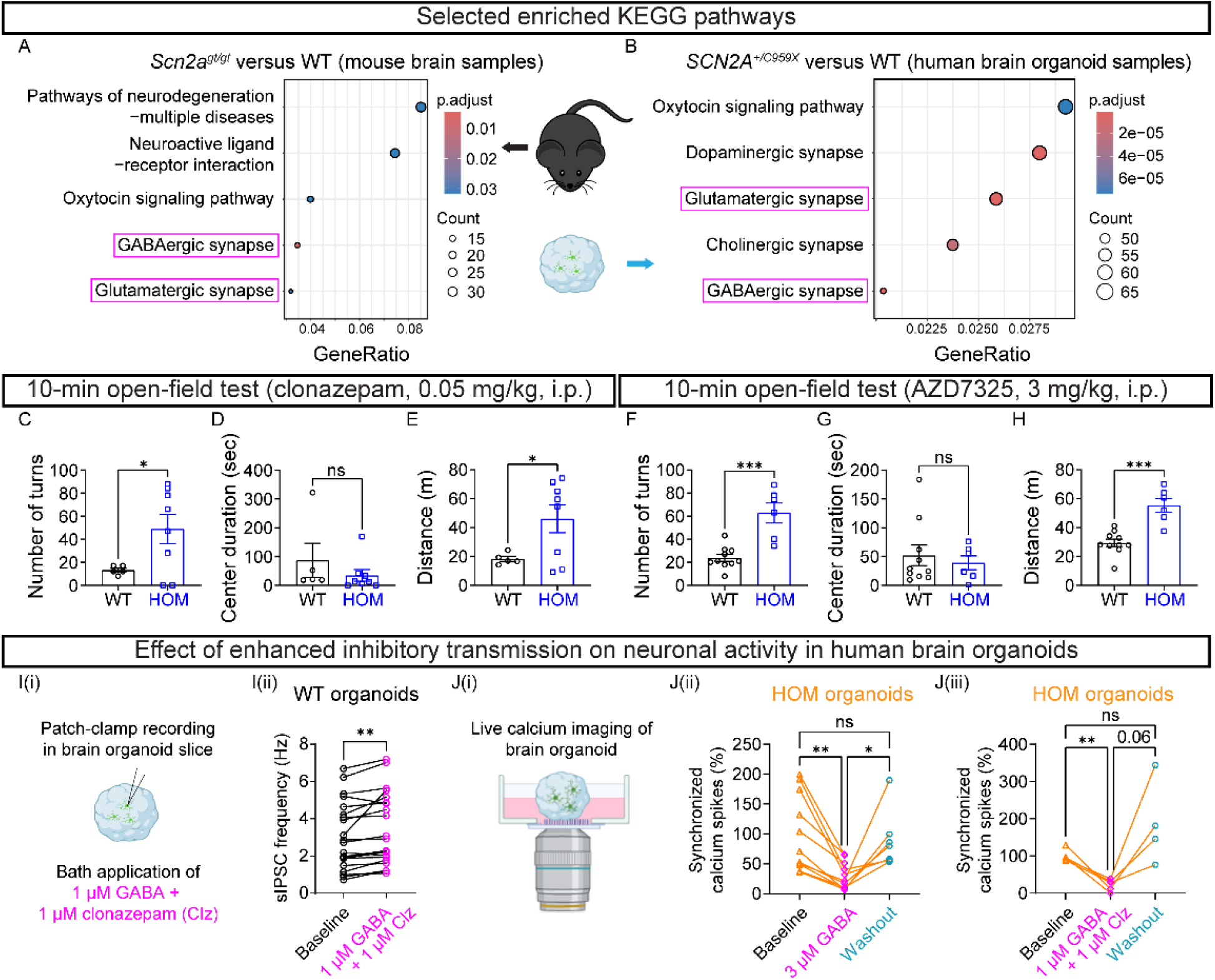
Acute effect of GABA_A_ PAMs on mice and human brain organoids. Related to Figure 5. (A, B) Selected enriched Kyoto Encyclopedia of Genes and Genomes (KEGG) pathways comparing *Scn2a^gt/gt^* (HOM) versus *Scn2a^+/+^*(WT) mice, and *SCN2A^+/C959X^* (HET) versus *SCN2A^+/+^*(WT) human brain organoids, respectively. GeneRatio represents the proportion of genes associated with each pathway, with dot size reflecting the gene count and color indicating the adjusted p-value (p.adjust). Notably, the glutamatergic synapse and GABAergic synapse pathways are enriched in both the mouse and human brain organoid models. (C–E) Open-field test following acute systemic administration of clonazepam (0.05 mg/kg, i.p.) showing number of turns (A), center duration (B), and distance traveled (C). n = 5 WT and n = 8 HOM mice. Unpaired Kolmogorov-Smirnov test (A), unpaired Mann-Whitney *U* test (B), and unpaired t test (C): ns, not significant; *p < 0.05. (F–H) Open-field test following acute systemic administration of AZD7325 (3 mg/kg, i.p.) showing number of turns (A), center duration (B), and distance traveled (C). n = 10 WT and n = 6 HOM mice. Unpaired Mann-Whitney *U* test (D) and unpaired t test (E, F): ns, not significant; ***p < 0.001. (I(i)) Schematic of patch-clamp recordings in brain organoid slices, and (I(ii)) effect of co-application of GABA (1 µM) and clonazepam (Clz, 1 µM) on sIPSC frequency in WT organoids before and after co-application of GABA (1 µM) and Clz (1 µM). n = 21 cells from 8 WT organoids. Paired t test: **p < 0.01. (J) Schematic of experimental setup for calcium imaging (J(i)) and pharmacological intervention in organoids. (J(ii)) Quantification of normalized synchronized calcium transient rates in *SCN2A^C959X/C959X^* (HOM) (n = 11 cells from 5 organoids for 3 μM GABA group) organoids before and after treatment with GABA. Mixed-effects analysis with Tukey’s multiple comparisons: ns, not significant; *p < 0.05; **p < 0.01. (J(iii)) Normalized calcium spike rates in HOM (n = 5 cells from 3 organoids) organoids under baseline, co-application of GABA (1 µM) and Clz (1 µM), and washout conditions. Mixed-effects analysis with uncorrected Fisher’s LSD: ns, not significant; **p < 0.01. Data are represented as mean ± SEM.

## RESOURCE AVAILABILITY

### Lead Contact

Requests for reagents and resources should be directed to the Lead Contact, Yang Yang.

### Materials Availability

This study did not generate new unique reagents.

### Data and Code Availability

Detailed datasets and source code supporting the current study are available from the corresponding author on request.

## EXPERIMENTAL MODEL AND SUBJECT DETAILS

### Mouse Strains

C57BL/6N-*Scn2a1^tm1aNarl^*/Narl (referred to as *Scn2a^+/gt^*) mice were generated from the National Laboratory Animal Center Rodent Model Resource Center based on a modified gene-trap (gt) design^21,22^. The generation and initial characterization of this mouse model have been previously described in our previous study^12,20^. All animal experiments were conducted in compliance with the Institutional Animal Care and Use Committee (IACUC) guidelines and the National Institutes of Health *Guide for the Care and Use of Laboratory Animals*. Mice were housed on a 12-hour light/dark cycle with *ad libitum* access to food (2018S Teklad, Envigo) and reverse osmosis water. Heterozygous (*Scn2a^+/gt^*) mice were used as breeding pairs to produce homozygous (*Scn2a^gt/gt^*) and wild-type (WT) littermates for experiments. Littermates of the same sex, but of different genotypes, were co-housed in ventilated cage racks containing 1/8” Bed-o’Cobb bedding (Anderson, Maumee, OH, USA) and provided with >8 g of enrichment material (e.g., shredded paper, crinkle-cut paper, or cotton nestlets). Genotyping was performed using ear-clipping samples. Behavioral experiments were conducted during the dark phase of the light/dark cycle at the Purdue Animal Behavioral Core. Mice of both sexes (3–5 months old) were acclimated to the behavioral testing room for 45–60 minutes prior to testing. All equipment was cleaned with a 70% isopropanol/water solution before and after each use. Animals were randomly assigned to treatment groups, and no animals were excluded from analysis.

### Reagents

3 mg/kg AZD7325 (IUPAC: 4-amino-8-(2-fluoro-6-methoxyphenyl)-N-propylcinnoline-3-carboxamide, MCE), 0.05 mg/kg clonazepam (IUPAC: 5-(2-chlorophenyl)-7-nitro-1,3-dihydro-1,4-benzodiazepin-2-one, Sigma-Aldrich), or 3 mg/kg CNO (Clozapine N-oxide dihydrochloride, Tocris) was administered intraperitoneally (i.p.) before behavioral testing.

### Antibodies

Primary antibodies used were Rabbit anti-SCN2A (Na_V_1.2) (1: 1000, Alomone Labs, ASC-002) and mouse anti-β-Actin (1:2000, Cell Signaling Technology, 3700S). Secondary antibodies were IRDye® 680RD Goat anti-Rabbit IgG Secondary Antibody (1:5000, LI-COR Biosciences, AB_10956166) and IRDye® 680RD Goat anti-Mouse IgG Secondary Antibody (1:5000, LI-COR Biosciences, AB_10956588).

## METHOD DETAILS

### Genotyping

Mice were labeled and genotyped via ear punch at weaning (21–28 days old). DNA was extracted from ear tissue using a tissue DNA extraction kit (Macherey-Nagel, Bethlehem, PA, USA). Genotyping for the trapping cassette was conducted using gene-specific PCR with the following primers: forward (5’ to 3’): GAGGCAAAGAATCTGTACTGTGGGG; reverse (5’ to 3’): GACGCCTGTGAATAAAACCAAGGAA. The PCR products were size-specific, with the WT allele generating a 240-base pair (bp) product and the gt allele producing a 340-bp product.

### Behavioral Assessments

#### Three-chamber social test (novel versus empty, novel versus toy)

Social preference was assessed using a standard three-chamber test (40.5 cm × 60 cm × 22 cm, Maze Engineers). Mice were given a choice between 1) an empty cylinder and a cylinder containing a same-sex novel mouse of similar age and size; and 2) a toy mouse and a novel mouse. Behavior was recorded for 5 minutes using EthoVision XT (Noldus).

#### Stranger and reciprocal interaction

Mice were placed in a clean cage with a novel mouse of the same sex, age, and size. Two paradigms were tested: 1) stranger interaction: pairing with a WT mouse; 2) reciprocal interaction: pairing with a mouse of the same genotype. Social behaviors (e.g., sniffing, following, rearing towards the novel mouse) were scored manually by three blinded researchers over a 10-minute session.

#### Resident-intruder test

Male mice were singly housed for six days without bedding changes to establish territoriality. On the seventh day, a male WT intruder mouse of similar age and size was introduced for 10 minutes. Interactions were recorded and scored for: 1) social behaviors: sniffing and following; 2) offensive behaviors: advancing, rearing towards, and fighting; and 3) defensive behaviors: retreating, rearing at the wall, and jumping away. Scoring was performed manually by three blinded researchers.

#### Home cage socialization

Home cage activity of mixed-genotype littermates of the same sex was recorded over 24 hours. Every 15 minutes, videos were assessed for social behaviors (e.g., sniffing, grooming, and physical contact) during both light and dark cycles.

#### Open field test

Mice were placed in an open field arena (40 cm × 40 cm × 40 cm, Maze Engineers) at 60 lux for 10 minutes. The center zone was defined as 20 cm square. EthoVision XT (Noldus) recorded: 1) total distance traveled; 2) time spent in the center zone; and 3) turning behavior. Each mouse participated only once.

#### Marble-burying test

Fifteen black glass marbles (1.5 cm diameter) were arranged in a 3 × 5 grid in a clean cage with 5 cm-deep bedding. Mice were allowed to interact with the marbles for 30 minutes. The number of marbles buried at least two-thirds was recorded.

#### Nesting test

Nesting height and quality were assessed as described previously^12^. Mice were singly housed in clean cages at the start of the dark cycle with one 5 cm × 5 cm square of compressed cotton nesting material (Ancare). 1) Height was recorded as the average of the four corners of the nest. 2) Quality was rated on a 0–5 scale (0 = untouched nesting material, 5 = well-formed cocoon-shaped nest).

### Adeno-Associated Virus (AAV) Production

pAAV-EF1a-mCherry-IRES-Flpo was a gift from Karl Deisseroth^66^ (Addgene plasmid # 55634 and viral prep # 55634-AAVrg; http://n2t.net/addgene:55634; RRID: Addgene_55634), AAV9-PHP.eB-EF1a-mCherry-IRES-Flpo with the titer of 2.56×10^13^ genome copies (GC)/mL was packed by Penn Vector Core (http://pennvectorcore.med.upenn.edu/); Control virus PHP.eB-Ef1a-DO-mCherry-WPRE-pA with the titer of 1.2×10^13^ GC/mL was packed by Bio-Detail Corporation^20^; pAAV-hSyn-mCherry was a gift from Karl Deisseroth (Addgene viral prep # 114472-AAV8; http://n2t.net/addgene:114472 ; RRID:Addgene_114472); pAAV-hSyn-hM3D(Gq)-mCherry was a gift from Bryan Roth (Addgene viral prep # 50474-AAV8; http://n2t.net/addgene:50474 ; RRID:Addgene_50474); pAAV-hSyn-hM4D(Gi)-mCherry was a gift from Bryan Roth (Addgene viral prep # 50475-AAV8; http://n2t.net/addgene:50475; RRID:Addgene_50475); pAAV.Syn.GCaMP6s.WPRE.SV40 was a gift from Douglas Kim & GENIE Project^67^ (Addgene viral prep # 100843-AAV1; http://n2t.net/addgene:100843; RRID:Addgene_100843); pENN.AAV.hSyn.HI.eGFP-Cre.WPRE.SV40 was a gift from James M. Wilson^68^ (Addgene viral prep # 105540-PHPeB; http://n2t.net/addgene:105540; RRID:Addgene_105540); pGP-AAV-syn-jGCaMP8m-WPRE was a gift from GENIE Project^69^ (Addgene viral prep # 162375-AAV1; http://n2t.net/addgene:162375; RRID:Addgene_162375).

### Surgical Procedures

For all surgeries (unless otherwise specified), mice were anesthetized systemically using a ketamine-xylazine cocktail and administered buprenorphine for postoperative analgesia and recovery support.

### AAV Injections

For systemic viral delivery, each adult mouse received either 2×10^11^ (low dose) or 4×10^11^ (high dose) viral particles of Flp or control AAV via tail vein injection. For brain injections, mice were anesthetized with a ketamine/xylazine cocktail (100/10 mg/kg, i.p.) and secured in a stereotaxic apparatus with ear bars (68046, RWD Ltd, China). A small midline scalp incision was made to expose the skull, and the periosteum was removed using 3% hydrogen peroxide. Small craniotomies (∼0.5 mm) were drilled over target sites (coordinates of the injection sites relative to bregma). Guide cannulas (62004, RWD Ltd, China) were inserted into the target brain regions at a controlled speed of 120 μm/min). AAVs (diluted to ∼5×10^12^ GC/mL in PBS containing 0.001% Pluronic F-68) were bilaterally injected under the following conditions: 1) control AAV or PHP.eB-Flp: AP +1.35 mm, ML ±1.00 mm, DV -3.30 mm and -4.50 mm; AP +0.50 mm, ML ±2.00 mm, DV -3.25 mm, 0.5–1 μL per site, 6 sites total; 2) AAVrg-Flp: (AP +1.35 mm, ML ±1.00 mm, DV -3.30 mm & −4.50 mm; 0.5–1 μL per site, 4 sites total. Injections were performed using a 33G injector (62204, RWD Ltd, China) connected to PEDAX tubing (62320, RWD Ltd, China) and a microliter syringe mounted on a microinjection syringe pump (World Precision Instruments, UMP3T-2). The tubing was pre-filled with water, followed by a layer of mineral oil (M5904, Sigma-Aldrich, USA) to separate the viral suspension. Viruses were injected at 100–150 nL/min, and the injector was left in place for 5–10 minutes post-injection before withdrawal. The incision was sutured, and mice were allowed to recover for one week, during which their health and body weight were closely monitored. Viral expression was confirmed *post hoc* via imaging of 50-μm brain sections containing the injection sites. Behavioral assessments and electrophysiological recordings were conducted after at least three weeks of viral expression.

For *in vivo* calcium imaging, mice received the following injections: 1) control AAV or AAVrg-Flp: AP +1.35 mm, ML +1.00 mm, DV -3.30 mm and -4.50 mm; AP +0.50 mm, ML ±2.00 mm, DV -3.25 mm, 1 μL per site, 4 sites total); 2) AAV cocktail (a 1:1 v/v mix of control AAV:AAV1-GCaMP6s or AAVrg-Flp:AAV1-GCaMP6s) into the right hemisphere striatum: AP +1.35 mm, ML -0.75 and -1.35 mm, DV -4.50 mm and -4.10 mm and -3.30 mm, 0.6 μL mix per site, 6 sites total. After the first round of calcium imaging, mice were administered 4×10^11^ viral particles of PHP.eB-Flp or PHP.eB-Cre via tail-vein injection for comparative “before-and-after” analysis.

### Perfusion and Tissue Processing

For immunostaining, mice were deeply anesthetized and transcardially perfused with ice-cold PBS, followed by 4% paraformaldehyde (PFA). For *LacZ* staining, 4% PFA was substituted with a solution of 2% formaldehyde and 0.2% glutaraldehyde in PBS (this substitution applies throughout this protocol). After perfusion, brains were carefully dissected and post-fixed overnight at 4°C in 4% PFA. Tissues were cryoprotected by sequential immersion in sucrose gradients (10%, 20%, and 30% sucrose in 0.01 M PBS) at 4°C until the samples fully sank. Subsequently, tissues were frozen in 20% and 30% sucrose solutions in 1× PBS for 24–48 hours before embedding. Samples were embedded in the Optimal Cutting Temperature (OCT) compound and frozen using dry ice, then stored at -80°C. Cryosections were cut at a thickness of 20 μm using a cryostat (Leica CM1950) and mounted onto slides. Sections were air-dried before analysis with a confocal microscope (Zeiss LSM 900).

### *LacZ* (β-galactosidase) Staining

Cryosections from *Scn2a^gt/gt^* and WT mice, with or without AAV injections, were processed simultaneously to minimize variation. Sections were fixed in 2% formaldehyde and 0.2% glutaraldehyde in PBS for 5 minutes at room temperature, then washed in PBS with 0.02% Triton X-100 for 5 minutes to reduce nonspecific antibody binding. Freshly prepared staining solution (X-Gal in Iron Buffer, 1:19, v/v) was applied to fully cover the tissue (∼50 μL per section) and incubated at 37°C in a humid chamber for 15–30 minutes, or longer if needed, until cells stained blue. Color development was monitored under a microscope. After staining, sections were washed three times with PBS, mounted in glycerol, and stored. Images were analyzed using an upright light microscope (Olympus).

### Immunostaining and Imaging Analysis

Cryosections (20 μm in thickness) were permeabilized, incubated in blocking buffer (0.5% Triton X-100 and 5% normal goat serum in PBS) for one hour at room temperature, and overlaid with primary antibodies overnight at 4°C. Then, the corresponding Alexa Fluor 488-, 594- or 647-conjugated secondary antibodies were applied. All stained sections were mounted with DAPI-containing mounting solution and sealed with glass coverslips. All immunofluorescence-labeled images were acquired using a confocal microscope^70^.

### RNA Sequencing

#### RNA extraction

Four *Scn2a^gt/gt^*(HOM) and four WT littermate mice were used to extract RNA. Mice were given an overdose of anesthesia and transcardiacally perfused with ice-cold PBS. Acute coronal brain slices (300-μm in thickness) were cut using a vibratome (Leica VT1200S, Germany). Tissues were rapidly microdissected, immersed into liquid nitrogen, and stored at -80°C until use (same procedures for Western Blotting and qPCR). Based on the manufacturer’s instructions, total RNAs were extracted with TRIzol reagent (Thermo Fisher Scientific, 15596018) from mouse cerebral tissues.

#### Library preparation and sequencing

Novogene prepared libraries using the TruSeq Stranded kit (Illumina, San Diego, CA), and RNA quality was assessed using an Agilent Nano RNA ChIP. Paired-end 150 bp reads were sequenced using the NovaSeq 6000.

#### Analysis

Reads were quality trimmed and Illumina TruSeq adapter sequences were removed using Trimmomatic v.0.36^71^. A sliding window approach to trimming was performed, using a window size of 5 and a required average Phred (quality) score of 16. Bases falling below a Phred score of 10 at the start and end of reads were trimmed and reads shorter than 20 bases in length after trimming were removed. FastQC v. 0.11.7^72^ was run to observe data quality before and after trimming/adapter removal. STAR v. 2.5.4b^73^ was used to align reads to the Ensembl *Mus musculus* genome database version GRCm38.p6. The htseq-count script in HTSeq v.0.7.0^74^ was run to count the number of reads mapping to each gene. HTSeq used Biopython v.2.7.3 in the analysis. HTSeq was run utilizing the GTF file on “intersection-nonempty” mode. The HTSeq feature was set to “exon” and the attribute parameter was set to “gene_id” and the --stranded=reverse option was set. The Bioconductor packages DESeq2 v.1.22.2^75^ and edgeR 3.24.3^76^ were used for differential expression analysis. Genes that were identified as differentially expressed in both packages were used as high-confidence differentially expressed genes and were used in subsequent pathway analysis. The Benjamini-Hochberg false discovery rate correction was used to correct p-values for multiple testing. To improve power, low-expression transcripts were filtered out of the data before performing differential expression analysis. The threshold chosen was to filter out all genes expressed at lower than 0.5 counts per million (CPM) in all combined samples. After filtering, 18,134 genes were remaining. The expression of genes between WT and HOM was deemed significant if the adjusted p-value < 0.05. The Bioconductor package biomaRt v. 2.38.0 was used to perform annotation of genes^77,78^. ClusterProfiler v. 3.10.1 was used to perform pathway and gene ontology enrichment analysis^79^.

### Western Blotting

Brain tissues were homogenized in ice-cold RIPA buffer (Thermo Fisher, 89901) with protease and phosphatase inhibitors (Thermo Fisher, A32953), sonicated, and centrifuged at 10,000× g for 10 minutes at 4°C. Protein concentrations were measured using a NanoDrop spectrophotometer (Thermo Scientific). Proteins were denatured in 1× sample buffer [62.5 mM Tris-HCl (pH 6.8), 2% SDS, 5% glycerol, 0.05% bromophenol blue] by boiling at 95°C for 5 minutes. Forty micrograms of total protein per sample were resolved on 8% SDS-PAGE gels and transferred onto PVDF membranes (Millipore, IPFL00010). Membranes were blocked with 5% nonfat milk in TBST (Tris-buffered saline with 0.1% Tween 20) for 1 hour at room temperature and incubated overnight at 4°C with primary antibodies diluted in 5% milk-TBST. Blots were washed three times in TBST (15 minutes each), incubated with IRDye® 680RD secondary antibodies (LI-COR Biosciences) for 2 hours at room temperature, and washed again (three 15-minute cycles). Immunoreactive bands were visualized using the Odyssey® CLx Imaging System (LI-COR Biosciences) and analyzed with Image Studio 6.0 (LI-COR Biotechnology) or FiJi. Protein levels were normalized to β-actin or GAPDH and further compared to WT littermate controls.

### Patch-clamp Recordings

#### Acute slice preparations

Slices were prepared from 3–5-month-old *Scn2a^gt/gt^* and control mice. Mice were deeply anesthetized with ketamine/xylazine (100/10 mg/kg, i.p., 0.1 mL/10 g body weight) before transcardial perfusion and decapitation. Brains were rapidly dissected into ice-cold slicing solution containing (in mM): 110 choline chloride, 2.5 KCl, 1.25 NaH_2_PO_4_, 25 NaHCO_3_, 0.5 CaCl_2_, 7 MgCl_2_, 25 glucose, 1 sodium ascorbate and 3.1 sodium pyruvate (pH 7.4, 305–315 mOsm, bubbled with 95% O_2_ and 5% CO_2_). Coronal slices (300 μm thick) containing the striatum were prepared using a vibratome (Leica VT1200 S, Germany). Slices were incubated for 10 minutes at 33°C in slicing solution, then transferred to artificial cerebrospinal fluid (aCSF; in mM; 125 NaCl, 2.5 KCl, 1.25 NaH_2_PO_4_, 25 NaHCO_3_, 2.0 CaCl_2_, 2.0 MgCl_2_, 10 glucose; pH 7.4, 305–315 mOsm, bubbled with 95% O_2_ and 5% CO_2_) for 10–20 minutes at 33°C before storage at room temperature for at least 30 minutes prior to recording.

#### Ex vivo whole-cell electrophysiology

Slices were placed in a recording chamber continuously perfused with aCSF at 32–33°C (2–3 mL/min). Neurons were visualized with an IR-DIC microscope (Olympus BX-51WI) equipped with an IR-2000 camera (Dage-MTI). Whole-cell patch-clamp recordings were performed on striatal medium spiny neurons (MSNs) identified by their characteristic morphology, including medium-sized, polygonal/dendritic spiny somata, and hyperpolarized resting membrane potentials (RMP < -80 mV)^20^. Thin-wall borosilicate pipettes (BF150-110-10, Sutter Instruments) with an open-tip resistance of 3–5 MΩ were fabricated using a P-1000 puller (Sutter Instruments).

Voltage-clamp recordings of spontaneous excitatory postsynaptic currents (sEPSCs), spontaneous inhibitory postsynaptic currents (sIPSCs), miniature excitatory postsynaptic currents (mEPSCs), and miniature inhibitory postsynaptic currents (mIPSCs) were performed to evaluate the excitatory/inhibitory (E/I) balance. The internal solution for these recordings consisted of (in mM): 120 CsMeSO_3_, 4 MgCl_2_, 0.2 EGTA, 10 HEPES, 4 Na_2_ATP, 0.3 Tris_3_-GTP, 14 Tris_2_-phosphocreatine, adjusted to pH 7.25 with CsOH (295–305 mOsm). Membrane potentials were held at *V*_hold_ = -80 mV for sEPSCs and mEPSCs and *V*_hold_ = 0 mV for sIPSCs and mIPSCs. Recordings of mEPSCs and mIPSCs were conducted in the presence of tetrodotoxin (TTX, 0.5 μM) to block action potential-driven activity, isolating spontaneous synaptic vesicle release. sEPSCs and sIPSCs reflected action potential-evoked transmitter release and ongoing synaptic activity. Each current type was recorded for 3 minutes in gap-free mode. Cells were included in the analysis if their leak current was <100 pA at *V*_hold_ = -80 mV. The sEPSC/sIPSC and mEPSC/mIPSC ratios were calculated for each neuron and averaged across cells^80^.

To confirm reductions in glutamate and GABA transmission, additional mEPSC and mIPSC recordings were conducted at *V*_hold_ = -80 mV using modified internal solutions. For mEPSCs, the internal solution included (in mM): 122 KMeSO_4_, 4 KCl, 2 MgCl_2_, 0.2 EGTA, 10 HEPES, 4 Na_2_ATP, 0.3 Tris-GTP, 14 Tris-phosphocreatine, adjusted to pH 7.25 with KOH (295– 305 mOsm). TTX (0.5 μM), picrotoxin (50 μM), and CGP55845 (2 μM) were used to block inhibitory synaptic transmission. For mIPSCs, the internal solution contained (in mM): 110 CsCl, 1 CaCl_2_, 5 MgCl_2_, 10 EGTA, 10 HEPES, 4 Na_2_ATP, 0.3 Tris-GTP, 14 Tris-phosphocreatine, adjusted to pH 7.25 with KOH (295–305 mOsm). TTX (0.5 μM), AP5 (50 μM), and CNQX (10 μM) were applied to block excitatory synaptic transmission.

Recordings were performed using an Axon MultiClamp 700B amplifier (Molecular Devices), and data were acquired with pClamp 11.4 software, filtered at 2 kHz, and sampled at 33 kHz with an Axon Digidata 1550B plus HumSilencer digitizer (Molecular Devices). Series resistance (Rs) was maintained at 15–30 MΩ, and recordings with unstable Rs (>20%) were excluded. Data files were saved in ABF 1.8 format and analyzed using the Mini Analysis Program (v6.08, Synaptosoft).

The internal solution for cell labeling included 0.1–0.2% neurobiotin. After recordings (∼30 minutes), slices were fixed in 4% paraformaldehyde (pH 7.4) for 20–30 minutes at room temperature, washed in PBS, and incubated overnight at 4°C with Alexa 488-conjugated streptavidin (1:250 in blocking solution)^81^.

### Neuropixels Recordings

#### Surgeries

Animal preparation was performed as described previously^33,34,82^. In brief, mice were anesthetized and head-fixed in a stereotaxic apparatus. The scalp and periosteum were removed, and a 0.5-mm craniotomy was drilled over the right striatum. The craniotomy was sealed with a circular piece of polydimethylsiloxane (PDMS) silicone. A metal headframe with a 10-mm circular opening (Narishige, MAG-1, and CP2) was attached to the skull using cyanoacrylate (Krazy Glue) and coated with Stoelting™ dental cement (Fisher Scientific). The cement formed a chamber to contain ground wire and aCSF during recordings. The craniotomy was covered with Kwik-Cast silicone sealant (World Precision Instruments). After a one-week recovery period, mice were treated with Rimadyl (2 mg/tablet, Bio-Serv) and handled daily to habituate them to the experimental rig over two weeks before recordings.

#### In vivo recordings

On the day of recording, mice were lightly anesthetized with isoflurane and head-fixed in the recording apparatus. A ground wire was secured to the skull, and the exposed brain was covered with 4% agar dissolved in aCSF. Before insertion, the Neuropixels probe was coated with CM-DiI for later identification. CM-DiI (Thermo Fisher Scientific, C7000) was dissolved in ethanol (1 μg/μL), applied to the probe tip, and dried. The Neuropixels probe was slowly inserted into the striatum (coordinates: AP +1.30 mm, ML ±1.25 mm, depth 5.10 mm) through the craniotomy at a rate of 120–480 μm/min. The probe was retracted slightly (to 5.00 mm depth) and allowed to settle for 10 minutes before recording began. Recordings were conducted using the first 384 electrodes of the probe, covering approximately 3.8 mm. After recording, the probe was retracted, cleaned with Tergazyme (Alconox), and rinsed with distilled water. Probe placement was confirmed by detecting DiI fluorescence in sectioned brain tissue.

#### Data acquisition and analysis

Electrophysiological recordings were performed using Neuropixels 1.0 probes at a 30-kHz sampling rate, controlled through the Open Ephys GUI (https://open-ephys.github.io/gui-docs/User-Manual/Plugins/Neuropixels-PXI.html). A 300-Hz high-pass filter was integrated into the probes, and an additional 300-Hz offline high-pass filter (third-order Butterworth) was applied before spike sorting. Spike sorting was performed with Kilosort 2.0, and outputs were refined manually in PHY^83^. Manual refinement included merging or splitting clusters and marking non-neural clusters as “noise” based on waveform shapes and autocorrelogram patterns. Noise clusters were identified by criteria such as a peak-to-trough ratio < 0.99 or a recovery slope < 0. Waveforms were extracted and averaged for each unit. High-quality striatal single units were identified and classified into four putative striatal cell types following established criteria^34^: MSNs, fast-spiking interneurons (FSIs), tonically active neurons (TANs), and unidentified interneurons (UINs). Units with narrow waveforms (Kilosort template trough-to-peak duration ≤ 400 μs) and a high proportion (>10%) of long interspike intervals (ISIs > 2 s) were classified as UINs and the other units with narrow waveforms were classified as FSIs. Units showing post-spike suppression lasting >40 m were categorized as TANs, and the remaining units were classified as MSNs.

### *In Vivo* Calcium Imaging in Freely Moving Mice

#### GRIN lens implantation and data collection

Mice underwent GRIN lens implantation^84,85^ two weeks after AAV injection. Animals were anesthetized with a ketamine/xylazine cocktail and secured in a stereotaxic apparatus. A ProView™ Integrated Lens (1.0 mm × 4.0 mm, Inscopix) was implanted into the right striatum (coordinates: AP +1.35 mm, ML +1.05 mm, depth 4.00 mm) at a controlled speed of 2 μm/s. The lens was affixed to the skull using dental cement (Metabond S380, Parkell). After one month of recovery, mice underwent 30-minute habituation sessions over 3–5 days in a linear three-chamber apparatus^55^ to acclimate them to the imaging setup. During these sessions, the miniscope was attached to the implanted lens.

On the test day, calcium imaging was performed at 10 Hz using the nVista system (Inscopix). The LED power was set to 0.1–0.3 mW at the miniscope focal plane. Imaging data were synchronized with behavioral data collected through EthoVision XT software (Noldus) using a DAQ card (USB-201, Measurement Computing). Behavioral tracking included a 10-minute baseline period followed by 10 minutes of social interaction with a novel stimulus mouse. Social interaction metrics included time spent in the social zone and sniffing duration.

For data analysis, calcium signals were processed using the Inscopix Data Processing Software (IDPS) with the following preprocessing parameters: spatial downsampling (by a factor of 2), temporal downsampling (by a factor of 1), and spatial bandpass filtering (frequency range: 0.005–0.5 Hz). Cell identification was performed using CNMFe with a peak-to-noise ratio (PNR) of approximately 9–10 and a cell diameter of ∼9 pixels. This preprocessing ensured optimal identification of individual neurons, extraction of fluorescence traces, and deconvolution of fluorescence signals into neuronal activity. Post-experiment, lens placement was verified by identifying the insertion site in sectioned brain tissue.

#### Identification of behaviorally tuned neurons

Behaviorally tuned neurons associated with sniffing behavior were identified using custom MATLAB (MathWorks) scripts^62^. The analytical pipeline included the following steps: 1) for each neuron *n*, the similarity between its calcium trace (ΔF/F) *Cn* and the behavior vector (*B*) was calculated as the normalized inner product: 2*B*·*Cn*/(|*B*|^2^+|*Cn*|^2^). The similarity value ranged from 0 (completely different vectors) to 1 (identical vectors). 2) behavior epochs were randomly shuffled, and the similarity between the shuffled behavior vector and each calcium trace (*Cn*) was calculated. This process generated a chance similarity distribution for each neuron. 3) Based on their similarity scores, ON neurons were classified as actual similarity exceeded the 99.17th percentile of the chance similarity distribution, indicating significantly higher activity during annotated behavior epochs; OFF neurons were classified as actual similarity fell below the 0.83rd percentile, indicating significantly lower activity during annotated behavior epochs; Others were classified as Neutral neurons, indicating no significant modulation by the behavior. These classifications were applied to all active neurons for each mouse, identifying ON and OFF neurons associated with the annotated behaviors.

### Human Brain Striatal Organoids

#### hiPSC culture and neural spheroid formation^86^

Human induced pluripotent stem cells (hiPSCs, WT: Kolf2.1, B07, C03, A11; heterozygous *SCN2A-C959X*: A02, E04, F01; homozygous *SCN2A-C959X*: A03, D06, F03) were maintained feeder-free on Matrigel (Corning, #354230) in StemFlex medium (Thermo Fisher, #A3349401) with daily media changes and passaged every 4–5 days using Versene (Thermo Fisher, #15040066). hiPSCs were dissociated using Accutase (Thermo Fisher, #NC9839010) and seeded (∼10,000 cells/well) in ultralow-attachment 96-well plates (Corning, #CLS3474) in Essential 8 medium (Thermo Fisher, #A1517001) supplemented with 10 μM Y-27632 (Selleck, #S1049). After centrifugation (100g, 3 min) and 24 hours of incubation at 37 °C, spheroids were transferred to Essential 6 medium (Thermo Fisher, #A1516401) with 2.5 μM dorsomorphin (Sigma-Aldrich, #P5499) and 10 μM SB-431542 (R&D, #1614) for 5 days.

#### Differentiation and Maturation

On day 6, spheroids were transferred to ultralow-attachment 6-well plates (Corning, #3471) and cultured in neural medium (Neurobasal-A, B-27 minus vitamin A, GlutaMAX, penicillin-streptomycin; Thermo Fisher) supplemented with 2.5 μM IWP-2 (Selleck, #S7085) and 50 ng/ml Activin A (PeproTech, #120-14P) with daily media changes. From day 12, SR11237 (100 nM, Tocris, #3411) was added. On day 22, maturation medium was supplemented with 20 ng/ml BDNF (PeproTech, #450-02), 20 ng/ml NT-3 (PeproTech, #450-03), 200 μM ascorbic acid (Wako, #323-44822), 50 μM dibutyryl-cAMP (Santa Cruz, #sc-201567A), and 10 μM DHA (MilliporeSigma, #D2534). From day 42, 2.5 μM DAPT (Stemcell, #72082) was added, and from day 46, cultures were maintained in neural medium with B-27 Plus (Thermo Fisher, #A3582801), with media changes every 4–5 days.

#### Calcium imaging and analysis^87^

Organoids were transduced with pGP-AAV-syn-jGCaMP8m-WPRE (Addgene, #162375-AAV1) (1 µL of 5×10^12^ GC/mL per organoid per well in a 24-well plate). After 3 weeks, organoids were transferred to glass-bottom 24-well plates (Cellvis, #P24-0-N) in BrainPhys medium (STEMCELL, #5790). Imaging was performed on a Zeiss LSM 900 confocal microscope (10× objective) after a 15-minute equilibration at 37 °C and 5% CO_2_. Baseline calcium activity was recorded (5 minutes, 488 nm, 50 ms interval, 512 × 512 resolution), followed by imaging after GABA (1 µM or 3 µM) or clonazepam (1 µM) application, and a final 5-minute washout.

Raw calcium traces were extracted using Inscopix Data Processing 1.9.2 and subsequently analyzed in MATLAB. Fluorescence signals (ΔF/F_0_) were computed as: (F(t)−F_0_)/F_0_, where F_0_ represents the lowest 10% of the fluorescence signal for each cell. Calcium transients were identified when ΔF/F_0_ > 0.5. Network bursts, defined as synchronized calcium spikes, were identified from population-averaged calcium signals using the following criteria: 1) peak amplitude > 10× noise s.d.; 2) ≥50% of total cells active during a calcium transient; and 3) cells included in ≥70% of bursts.

#### RNA sequencing and differential gene expression analysis

##### RNA extraction, library preparation, and sequencing

RNA was extracted from 7 WT (Kolf2.1, B07, C03) and 7 heterozygous *SCN2A-C959X* (A02, E04, F01) 5-month-old organoids (RNeasy Mini Kit, QIAGEN, #74104). Poly(A)+ RNA (100–250 ng) was isolated using NEBNext® Poly(A) mRNA Magnetic Isolation (NEB) and fragmented on-bead using xGen RNA Library Kit (IDT). Libraries were sequenced on an Illumina NovaSeq X+ (300-cycle 25B flow cell, 30–36M paired-end reads/sample). Fastq files were generated using BCL2FASTQ (v1.8.4).

##### Data processing and alignment

Reads were quality-filtered using fastp (v0.23.2) (PMID: 30423086) (phred ≥30, read length ≥50 bp) and aligned to GRCh38 (Ensembl release 104) using STAR (v2.7.10a) (PMID: 23104886) in two-pass mode. *SCN2A* variant calling was performed using GATK HaplotyperCaller (v4.2.2.0) (PMID: 20644199) with joint-genotyping (PMID: 31249686) to confirm WT, HET, and HOM sample genotypes.

##### Gene expression and batch correction

Gene counts were assigned using featureCounts (v1.6.1) (PMID: 24227677) in paired-end, reverse-stranded mode. Initial clustering was assessed using DESeq2 (v1.34.0) (PMID: 25516281) in R (v4.1.3). Batch effects were computed using RUVSeq (v1.28.0) (PMID: 25150836). Genes with ≥5 reads in ≥2 samples were retained, and normalization was applied via the upper-quartile method with betweenLaneNormalization.

##### Differential expression and pathway analysis

Differential gene expressions were analyzed using edgeR (v3.36.0) (PMID: 19910308) with a quasi-likelihood negative binomial model, incorporating RUVSeq batch-correction factors (k = 5). Adjusted p-values were computed using Benjamini-Hochberg correction, and genes with FDR < 0.05 were considered differentially expressed (DEs). Functional enrichment for KEGG, Reactome, and Gene Ontology (GO) was performed using clusterProfiler (v4.10.0) (PMID: 22455463) in R (v4.3.2). Ingenuity Pathway Analysis (PMID: 24336805) was used for disease and pathway annotations.

##### Visualization

Batch-corrected counts per million (CPM) values were extracted from edgeR for heatmap generation (ComplexHeatmap (v2.14.0) (PMID: 38868715) in R (v4.2.1)). Volcano plots were visualized using EnhancedVolcano (v1.16.0)^88^.

## QUANTIFICATION AND STATISTICAL ANALYSIS

Normality and variance similarity were measured by GraphPad Prism before the application of any parametric tests. All data were analyzed blind to genotype. If no significant sex differences were observed, data from both sexes were combined for analysis. Behavior data were analyzed by repeated-measures two-way analysis of variance (ANOVA) with a Bonferroni *post-hoc* unless otherwise specified. Neuropixels data were analyzed using a linear mixed-effects model. For comparisons between two groups, either a two-tailed Student’s t test (parametric) or an unpaired two-tailed Mann-Whitney *U* test (non-parametric) was applied. For multiple comparisons, data were analyzed using one-way or two-way ANOVA with Tukey’s post-hoc correction (parametric) or Kruskal-Wallis test with Dunn’s multiple comparison correction (non-parametric), as appropriate. *Post hoc* tests were conducted only when the primary analysis showed statistical significance. Error bars in all figures represent the mean ± SEM. Statistical significance was defined as p < 0.05. Significance levels are denoted as follows: p < 0.05 is indicated as *, p < 0.01 is indicated as **, p < 0.001 is indicated as ***, and p < 0.0001 is indicated as ****. Mice from different litters and body weights were randomly stratified by sex into treatment groups. No additional randomization methods were applied in animal studies.

